# Parity-induced changes to mammary epithelial cells control NKT cell expansion and mammary oncogenesis

**DOI:** 10.1101/2021.08.23.457350

**Authors:** Amritha V. Hanasoge Somasundara, Matthew A. Moss, Mary J. Feigman, Chen Chen, Samantha L. Cyrill, Michael F. Ciccone, Marygrace C. Trousdell, Macy Vollbrecht, Siran Li, Jude Kendall, Semir Beyaz, John E. Wilkinson, Camila O. dos Santos

## Abstract

Pregnancy reprograms the epigenome of mammary epithelial cells (MECs) in a manner that control responses to pregnancy hormone re-exposure and the rate of carcinoma progression. However, the influence of pregnancy on the tissue microenvironment of the mammary gland is less clear. Here, we used single-cell RNA sequencing to comparatively profile the composition of epithelial and non-epithelial cells in mammary tissue from nulliparous and parous female mice. Our analysis revealed an expansion of γδ Natural Killer T (NKT) immune cells following pregnancy, in association with upregulation of immune signal molecules in post-pregnancy MECs. We show that expansion of NKT cells following pregnancy is due to elevated expression of the antigen presenting molecule CD1d protein, which is known to induce NKT activation. Accordingly, loss of CD1d expression on post-pregnancy MECs, or overall lack of activated NKT cells, accompanied the development of mammary oncogenesis in response to cMYC overexpression and loss of Brca1 function. Collectively, our findings illustrate how pregnancy-induced epigenetic changes modulate the communication between MECs and the mammary immune microenvironment, and establish a causal link between pregnancy, the immune microenvironment, and mammary oncogenesis.

## Introduction

Changes to the functions of immune cells modulate both the mammary immune microenvironment and mammary epithelial cells (MEC) lineages during all stages of mammary development. For example, CD4+ T-helper cells guide lineage commitment and differentiation of MECs, while macrophages provide growth factors and assist in removal of cellular debris arising from apoptotic events, during postnatal stages of mammary development (Dawson et al., 2020; Hitchcock et al., 2020; Plaks et al., 2015; Rahat et al., 2016; Stewart et al., 2019; Wang et al., 2020). Accordingly, changes that impact immune cell function and abundance can also influence the development and progression of mammary oncogenesis (Bach et al., 2021; Ibrahim et al., 2020).

Immune surveillance and communication in the mammary gland are critical to post-pregnancy mammary tissue homeostasis, particularly as part of mammary reconstruction during post-partum involution. Alterations to immune cell composition during mammary gland involution have also been suggested to influence mammary tumor progression (Lyons et al., 2011). For example, T-cell activity is suppressed by the infiltration of involution-associated macrophages, an immune reaction that may also induce mammary tumorigenic development (Martinson et al., 2015)(Freire-de-Lima et al., 2006)(Guo et al., 2017)(Fornetti et al., 2012) (O’Brien et al., 2010).

Conversely, cell-autonomous processes in MECs contribute to pregnancy-induced breast cancer protection, a life-long lasting effect that decreases the risk of breast cancer by ~30% in rodents and humans (Medina et al., 2004)(Britt et al., 2007)(Terry et al., 2018). For example, p53 function is critical for blocking mammary tumor development in murine and human MECs, with a complete loss of p53 in post-pregnancy MECs promoting tumorigenic initiation (Sivaraman et al., 2001)(Medina and Kittrell, 2003). Epigenetic-mediated alterations of post-pregnant MECs have been shown to interfere with the transcriptional output of cMYC, which suppressed mammary oncogenesis via oncogene-induced senescence (Feigman et al., 2020). Given that oncogene-induced senescence signals influence the immune system, a link between normal pregnancy-induced mammary development, the immune microenvironment, and oncogenesis needs to be addressed to fully understand the effects of pregnancy on breast cancer development.

In this study, we characterize the interactions between cell-autonomous (MECs) and non-cell-autonomous (immune cells) factors that occur as part of normal pregnancy-induced mammary development, and that are involved in repressing cancer development in the post-involuted mammary gland. Our analysis identified that pregnancy induces the expansion of a natural killer T-cell (NKT) population during the late stages of involution, which preferentially populates the fully involuted mammary tissue. Unlike the typical NKT cells that bear αβTCRs, mammary resident, pregnancy-induced NKT cells express γδTCRs on their surface, indicating a role in specialized antigen recognition. NKT cell expansion was linked with increased expression of the antigen-presenting molecule, CD1d, on the surface of post-pregnancy MECs, which was associated with the stable gain of active transcription markers at the Cd1d loci and increased mRNA levels. Further analysis demonstrated that gain of CD1d expression on post-pregnancy MECs, and expansion of γδNKT cells was observed in mammary tissues that failed to develop premalignant lesions and tumors in response to oncogenic signals, such as either cMYC overexpression or loss of Brca1, thus connecting pregnancy-induced molecular changes with alteration of immune microenvironment and lack of mammary oncogenesis. Altogether, our findings elucidate how signals brought to MECs during pregnancy-induced development regulate epigenomic changes, gene expression, and immune surveillance, which together control mammary oncogenesis.

## Results

### Single cell analysis identifies transcriptional programs and immune cellular heterogeneity in mammary tissue from parous female mice

The utilization of single cell strategies has elucidated the dynamics of epithelial cell lineage specification and differentiation across major mammary developmental stages (Bach et al., 2017; Chung et al., 2019; Li et al., 2020a; Pal et al., 2017, 2021). Previous studies have indicated that post-pregnancy epithelial cells bear an altered transcriptome and epigenome, thus suggesting that pregnancy stably alters the molecular state of this cell type (Blakely et al., 2006; Feigman et al., 2020; Huh et al., 2015; dos Santos et al., 2015) . However, it remains unclear whether pregnancy leads to disproportionate changes in the transcriptome of specific mammary cell populations, which we investigated in this study.

In order to characterize the effects of parity on the cellular composition and heterogeneity of mammary glands, we used single cell RNA-sequencing (scRNA-seq) to compare the abundance, identity and gene expression of mammary gland epithelial and non-epithelial cells from nulliparous (virgin, never pregnant) and parous female mice (20 days gestation, 21 days lactation, 40 days post-weaning). scRNA-seq clustering defined 20 cellular clusters (TCs), which were further classified into 3 main cell types; epithelial cells (Krt8+ and Krt5+), B-lymphocytes (CD20+), and T-lymphocytes (CD3e+), and 2 smaller clusters, encompassing fibroblast-like cells (Rsg5+) and myeloid-like cells (Itgax+), with similar cell cycle state **(Supplementary Fig. 1A-C).**

To characterize the cellular heterogeneity across pre- and post-pregnancy MECs, we used a re-clustering approach, which selected for cells expressing the epithelial markers Epcam, Krt8, Krt18, Krt14 and Krt5, and resolved 11 clusters of mammary epithelial cells (ECs) (Henry et al., 2021)(**Fig. 1A**). Analysis of cellular abundance and lineage identity revealed that clusters EC7 (mature myoepithelial MEC), EC9 (luminal common progenitor-like MEC), EC10 and EC11 (bi-potential-like MECs), were evenly represented in pre- and post-pregnancy mammary tissue, thus demonstrating populations of cells that are mostly unchanged by a pregnancy cycle. We also identified clusters predominantly represented within pre-pregnancy MECs (EC2, EC4, and EC8), and those biased towards a post-pregnancy state (EC1, EC3, EC5, and EC6), classified as luminal alveolar-like clusters (EC1, EC2 and EC6), myoepithelial progenitor-like clusters (EC3 and EC4), and luminal ductal-like clusters (EC5 and EC8) (**Supplementary Fig. S1D, S1E and S1F**). Comparative gene expression analysis indicated that processes associated with immune cell communication were markedly enriched in luminal and myoepithelial cell clusters biased towards the post-pregnancy state (**Fig.1B, Supplementary Fig. S2A-B and Supplementary File S1**). This observation was supported by analysis of previously published pre- and post-pregnancy bulk RNA-seq data, which suggested an overall enrichment for immune communication signatures in epithelial cells after a full pregnancy cycle (Feigman et al., 2020) (**Supplementary Fig. 2C and Supplementary File S2**).

**Figure 1.**
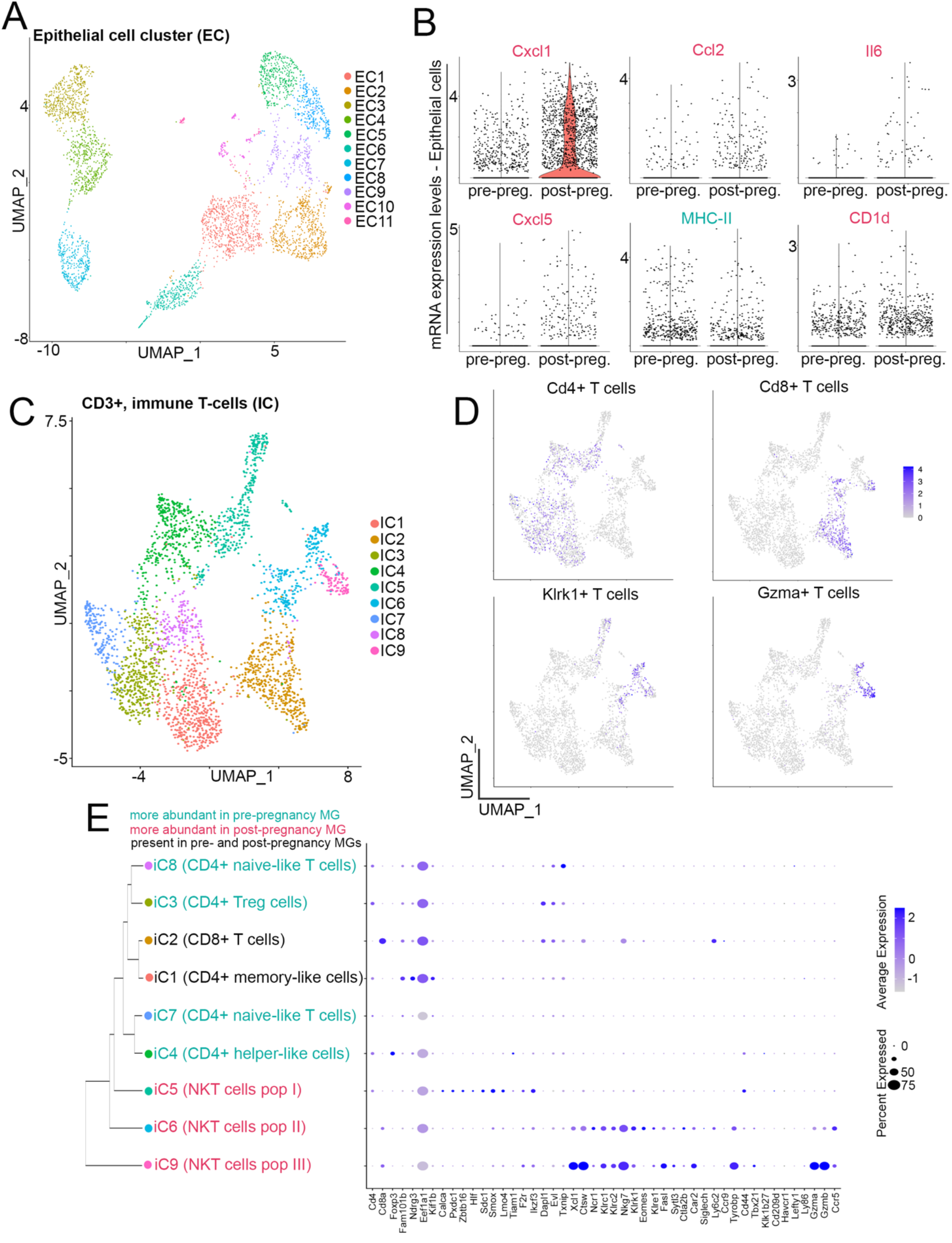
Single cell analysis identifies transcriptional programs and immune cellular heterogeneity in mammary tissue from parous female mice. (**A**) UMAP showing epithelial-focused re-clustering (Epcam+, Krt8+, Krt18+, and Krt5+ cells) of pre- and post-pregnancy MECs. (**B**) mRNA levels of senescence-associated, immune communication genes *Cxcl1*, *Ccl2*, *Il6*, *Cxcl5*, *Mhc-ii* and *Cd1d* in pre- and post-pregnancy MECs. (**C**) UMAP showing T-cell focused re-clustering (CD3e+ cells) of pre- and post-pregnancy mammary resident immune cells. (**D**) Feature plots showing the expression of T cell markers Cd4, Cd8, Klrk1 and Gzma. (**E**) Dendrogram clustering and dot plot showing molecular signature and lineage identity of pre- and post-pregnancy mammary resident CD3+ immune cells.

Changes in the immune microenvironment are known to contribute to pregnancy-induced mammary development (Coussens and Pollard, 2011). A series of single cell strategies have identified alterations to mammary immune composition across several stages of mammary gland development and cancer development (Bach et al., 2021; Dawson et al., 2020; Saeki et al., 2021). However, it still unclear whether the immune composition of fully involuted, post-pregnancy mammary tissue resembles the pre-pregnancy mammary state, or whether a combination of epithelial and non-epithelial signals collectively influence the normal and malignant development of mammary tissue. In light of the potentially altered epithelial-to-immune cell communication identified in post-pregnancy MECs suggested above, we set out to understand the effects of pregnancy on the mammary resident immune compartment using scRNA-seq. Transcriptional analysis of clusters representing B-lymphocytes (CD20+) did not identify major differences between cells from pre- or post-pregnancy mammary glands, suggesting that B-cells may not be significantly altered in fully involuted mammary tissue (**Supplementary Fig. S3A**). Re-clustering of CD3e+ T-lymphocytes identified 9 distinct immune cell clusters (IC) marked by the expression of immune lineage genes such as Cd4, Cd8, Klrk1, and Gzma (**Fig. 1C-D**). Interestingly, classification according to cell abundance and lineage identity of pre- and post-pregnancy mammary resident lymphocytes, revealed 2 cellular clusters, IC1 (CD4+ memory-like T-cells), and IC2 (CD8+ T-cells), which were evenly represented across pre- and post-pregnancy mammary tissue (**Supplementary Fig. S3B-C**). Differential gene expression analysis of pre- and post-pregnancy T cells classified under clusters IC1 and IC2 identified minimal expression changes, suggesting that the transcriptional output of CD8 T-cells (IC2), and certain populations of CD4+ T-cells (IC1) were not substantially altered by parity (**Supplementary Fig. S3D-E**).

Analysis of clusters biased towards pre-pregnancy mammary tissue identified several populations of CD4+ T-lymphocytes, with gene identifiers supporting their identity as CD4 Tregs (IC3), CD4+ naïve T-cells (IC7 and IC8), and CD4+ helper T-cells (IC4), suggesting pre-pregnancy mammary tissues are enriched for populations of CD4+ T-cells (**Fig. 1E**). Conversely, clusters enriched with post-pregnancy mammary immune cells (IC5, IC6, and IC9) were classified as NKT cells, a specialized population of T-cells involved in immune recruitment and cytotoxic activity (Godfrey et al., 2004) (**Fig. 1E**). Such clusters expressed master regulators of NKT cellular fate, including transcription factors (TFs) Tbx21 (Tbet), and Zbtb16 (Plzf) (Townsend et al., 2004) (Savage et al., 2008).

While Natural killer (NK) cells are known to play a role in mammary gland involution and parity-associated mammary tumorigenesis (Fornetti et al., 2012; Martinson et al., 2015), the role of NKT cells in this process has yet to be determined. Therefore, we set out to analyze clusters of immune cells expressing the common NK/NKT marker Nkg7 in order to further define the influence of pregnancy on the abundance and identity of NK and NKT cells. Deep-clustering analysis of Nkg7+ immune cells revealed 6 distinct cell clusters (NC1-6). Cells classified under cluster NC5, which includes cells from both the pre- and post-pregnancy mammary tissue, lacked expression of CD3e, and therefore was the only cluster with a NK cell identity in our dataset (**Supplementary Fig. S4A-C**). Further gene expression analysis confirmed that post-pregnancy mammary glands are enriched with a variety of NKT cells, including those expressing markers of cell activation (Gzmb and Ccr5) and of a resting state (Bcl11b) (**Supplementary Fig. S4C**). In agreement, each of the post-pregnancy-biased NKT cell clusters were enriched with an array of immune activation signatures, suggesting an altered state for these cell populations after pregnancy (**Supplementary Fig. S4D**).

Collectively, our scRNA-seq analysis of fully involuted mammary tissue confirmed that pregnancy leads to a stable alteration of the transcriptional output of post-pregnancy MECs, including gene expression signatures that suggest enhanced communication with the mammary immune microenvironment. In addition, our study indicates that mammary resident NKT cells are present at higher levels in post-pregnancy glands, further suggesting that pregnancy plays a role in inducing changes to the mammary immune microenvironment.

### Pregnancy induces the expansion of a specific population of NKT cells

During post-partum mammary gland involution, the immune composition of the gland expands with an influx of infiltrating mast cells, macrophages, neutrophils, dendritic cells and natural killer cells, which remove apoptotic epithelial cells and support the remodeling of the gland (Guo et al., 2017; Kordon and Coso, 2017; O’Brien et al., 2010; Schwertfeger et al., 2001). Since our scRNA-seq analyses suggested that fully involuted, post-pregnancy mammary glands are enriched for populations of NKT cells, we next utilized a series of flow cytometry analyses to validate this observation.

Analysis using antibodies against the markers NK1.1 and CD3, which defines NKT cells (NK1.1+CD3+), identified a 12-fold increase in the abundance of NKT cells in post-pregnancy mammary tissue, consistent with the results of our scRNA-seq data (**Fig. 2A**). Further analysis indicated a 2.3-fold higher abundance of NKT cells in recently involuted mammary tissue (15 days post offspring weaning), compared to mammary glands from nulliparous mice, or those exposed to pregnancy hormones for 12 days (mid-pregnancy), suggesting that the expansion of NKT cells is likely to initiate at the final stages of post-pregnancy mammary involution (**Supplementary Fig. S5A**). The selective expansion of NKT cells was further supported by the analysis of markers that define mammary resident neutrophils (Ly6G+), and mammary resident macrophages (CD206+), which were largely unchanged between pre- and post-pregnancy mammary tissue (**Supplementary Fig. S5B-C**). Immunofluorescence analysis of Cxcr6-GFP-KI mammary tissue, previously described to selectively label NKT cells (Germanov et al., 2008), demonstrated several GFP+ cells surrounding mammary ductal structures from pre-pregnancy mammary tissue, an observation that supports the presence of NKT cells in mammary tissue (**Supplementary Fig. S5D**). Moreover, analysis of bone marrow and spleen from nulliparous and parous mice showed no difference in the abundance of NK1.1+CD3+ cells, suggesting that the pregnancy-induced expansion of NKT cells is mammary-specific (**Supplementary Fig. S5E-F**).

**Figure 2.**
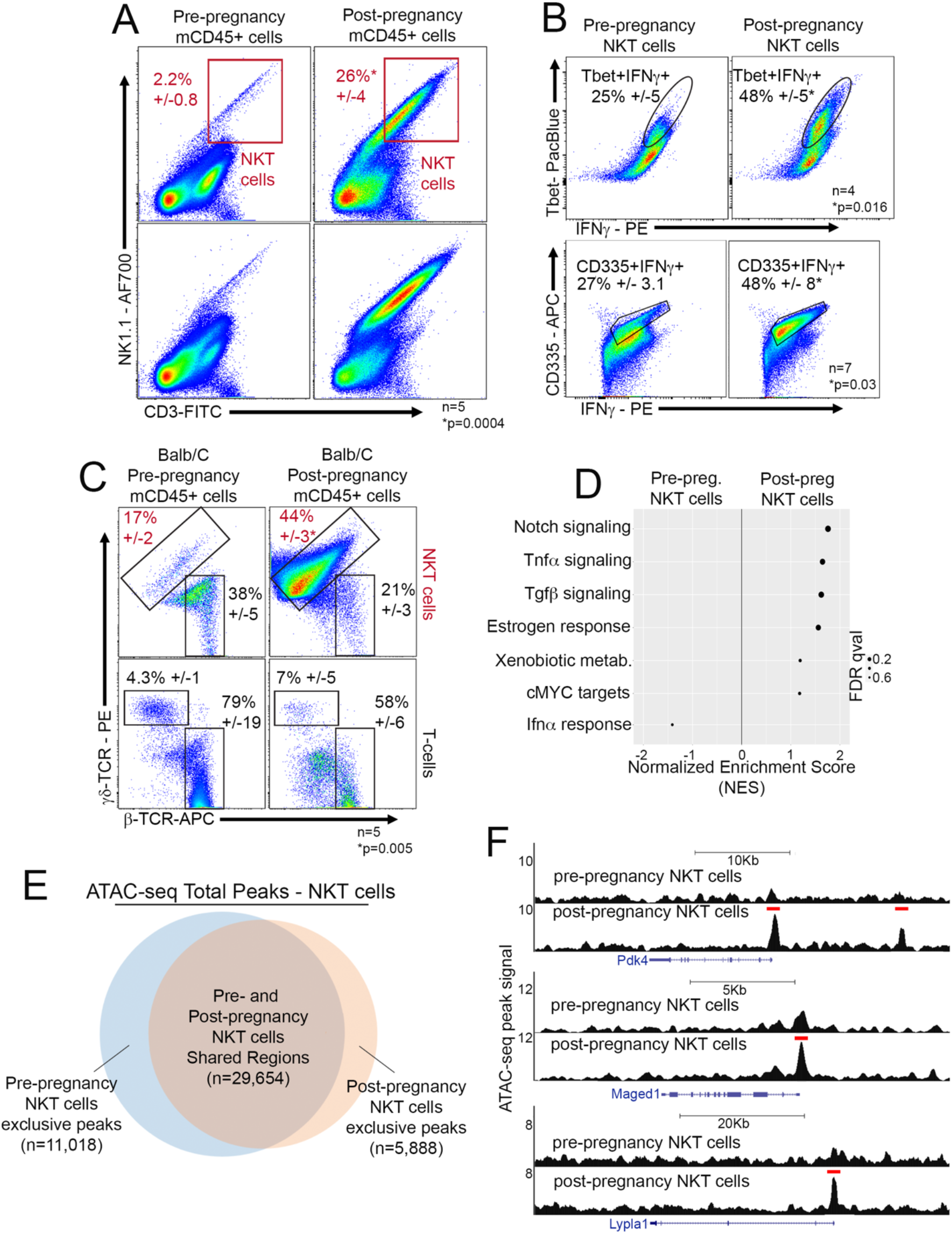
Pregnancy induces expansion of specific populations of NKT cells. (**A**) Flow cytometry analysis of resident CD45+ cells harvested from pre- and post-pregnancy mammary tissue, and their distribution of NKT cells (NK1.1+CD3+). n=5 nulliparous and 5 parous female mice. *p=0.0004. (**B**) Flow cytometry analysis of the classical NKT cell markers T-bet, CD335, and IFNγ in NKT cells harvested from pre- and post-pregnancy mammary tissue. For Tbet analysis n=4 nulliparous and 4 parous female mice. *p=0.016. For CD335 analysis n=7 nulliparous and 7 parous female mice. *p=0.03. (**C**) Flow cytometry analysis of β and γδ T-cell receptors (TCRs) of pre- and post-pregnancy mammary NKT cells. n=5 nulliparous and 5 parous female mice. *p=0.005. (**D**) Gene set enrichment analysis of differentially expressed genes in FACS-isolated NKT cells from pre- and post-pregnancy mammary tissue. (**E**) Venn-diagram demonstrating the number of shared and exclusive ATAC-seq peaks of FACS-isolated NKT cells from nulliparous female mice (blue circle) and parous female mice (orange circle). (**F**) Genome browser tracks showing distribution of MACS-called ATAC-seq peaks at the Pdk4, Maged1 and Lypla1 genomic loci from pre- and post-pregnancy NKT cells. For all analyses, error bars indicate standard error of mean across samples of the same experimental group. Statistically significant differences were considered with Student’s t-test *p*-value lower than 0.05 (p<0.05).

To further characterize the identity of the post-pregnancy, mammary resident NKT cells, we combined cell surface and intracellular staining to detect canonical NKT lineage markers, including the NKT master regulator Tbet, the NKT/T-cell secreted factor IFNγ, and the NKT lineage marker Nkp46 (CD335) (Yu et al., 2011). Pre- and post-pregnancy, mammary resident NK1.1+CD3+ cells expressed all three markers, supporting their NKT identity. However, we detected a 2-fold increase in the percentage of post-pregnancy cells expressing Tbet, IFNγ, and CD335, suggesting that specific populations of NKTs are expanded in post-involuted mammary tissue (**Fig. 2B**).

We also investigated whether pregnancy induced NKT cells represented a specialized population of CD8+ T-cells, a cytotoxic cell type recently reported to reside in mammary tissues (Wu et al., 2019). We found that a fraction of the NKT cells present in both pre- and post-pregnancy mammary tissue expressed CD8 on their surface, accounting for 41% and 35% of the total NKT cells, respectively (**Supplementary Fig. S5G**). To determine whether the triple-positive (CD3+NK1.1+CD8+) cells contributed significantly to the expanded population of post-pregnancy NKT cells, we analyzed mammary tissue of nulliparous and parous RAG1 KO mice, which lack mature CD8+ T-cells (Mombaerts et al., 1992). We observed a 10-fold expansion of NKT cells in RAG1 KO post-pregnancy mammary tissue, suggesting that CD8-expressing cells do not comprise a significant fraction of pregnancy-induced NKT cells (**Supplementary Fig. S5H**). These results are consistent with our scRNA-seq data, and further validate the existence of specific NKT subtypes in mammary glands after a full pregnancy cycle.

NKT cells have multiple roles, including tissue homeostasis, host protection, microbial pathogen clearance, and anti-cancer activity, mediated through their ability to recognize both foreign- and self-antigens via T-cell receptors (TCRs) (Balato et al., 2009). Therefore, we next investigated changes to the TCR repertoire of mammary resident, post-pregnancy NKT cells. We found that 17% of NKT cells expressed γδTCRs, in marked contrast to post-pregnancy NKT cells, which mostly expressed γδTCR chains (44%) (**Fig. 2C, top panel**). A pregnancy cycle did not alter TCR composition across all immune cells, given that mammary resident, pre- and post-pregnancy CD8+ T-cells mostly express αβTCRs, suggesting that parity promotes expansion of specific subtypes of NKT cells that bear a specific TCR repertoire (**Fig. 2C, bottom panel**).

We next investigated the molecular signatures of FACS-isolated, mammary resident, NKT cells. Unbiased pathway analysis of bulk RNA-seq datasets revealed the enrichment of post-pregnancy NKT cells for processes controlling overall NKT development and activation, such as Notch signaling, TNF α signaling, Tgfβ signaling, response to estrogen, and cMYC targets (Oh et al., 2015)(Almishri et al., 2016)(Doisne et al., 2009)(Huber, 2015)(Mycko et al., 2009). Conversely, pre-pregnancy NKT cells were mainly enriched for processes previously associated with reduced immune activation, such as IFNa response (Bochtler et al., 2008) (**Fig. 2D, Supplementary File S3**).

The activation of specific processes in post-pregnancy NKT cells was also evident from analysis of their accessible chromatin landscape. ATAC-seq profiles showed similar genomic distributions of accessible regions across pre- and post-pregnancy NKT cells, with a 93% overlap of their total accessible chromatin regions, suggesting that parity-induced changes did not substantially alter the chromatin accessibility associated with NKT lineage (**Fig. 2E and Supplementary Fig. S6A**). General TF motif analysis identified chromatin accessible regions bearing classical NKT regulator DNA binding motifs such as T-bet, Plzf, and Egr2, further supporting their NKT lineage identity (Seiler et al., 2012) (**Supplementary Fig. S6B**). Analysis of accessible chromatin exclusively to post-pregnancy NKT cells showed an enrichment for terms/genes associated with regulation of the adaptive immune response, killer cell activation and antigen presentation, such as Pdk4, Maged1, and Lypla1, all involved in enhanced immune-activation (Na et al., 2020)(Connaughton et al., 2010)(Lee et al., 2016)(Jehmlich et al., 2013) (**Fig. 2F and Supplementary Fig. S6C**). DNA motif analysis at accessible regions exclusive to post-pregnancy NKT cells identified enrichment of specific TF motifs, including those recognized by MAF, a factor associated with an activated NKT state, and previously predicted by our scRNA-seq data to be expressed in cell clusters with an NKT identity (**Supplementary Fig. S6D**).

Overall, our analysis confirmed that post-pregnancy mammary tissue has an altered γδNKT cell composition, which bears molecular and cellular signatures of activated and mature adaptive immune cells.

### NKT expansion requires CD1d expression on post-pregnancy MECs

Classically, NKT cells are subdivided based on their activating antigens, including the main antigen-presenting molecules MHC class I, MHC class II, and the non-classical class I molecule, CD1d, which can be expressed on the surface of macrophages and dendritic cells, and as well on the surface of epithelial cells (Gapin et al., 2013; Rizvi et al., 2015; Thibeault et al., 2009). Therefore, we next analyzed whether the expression of antigen-presenting factors on the surface of mammary epithelial and non-epithelial cells could underlie NKT cell expansion after pregnancy.

Flow cytometry analysis detected a 5-fold increase in the CD1d levels on the surface of post-pregnancy luminal and myoepithelial MECs (**Fig. 3A-B**). In contrast, no differences in the expression of antigen-presenting factors MHC-I and MHC-II on the surface of pre- and post-pregnancy MECs were found (**Supplementary Fig. S7A-B**). No difference in surface expression of CD1d on mammary CD45+ immune cells was detected, suggesting that signals provided by CD1d+ MECs could promote the post-pregnancy expansion of mammary NKT cells (**Supplementary Fig. S7C**).

**Figure 3.**
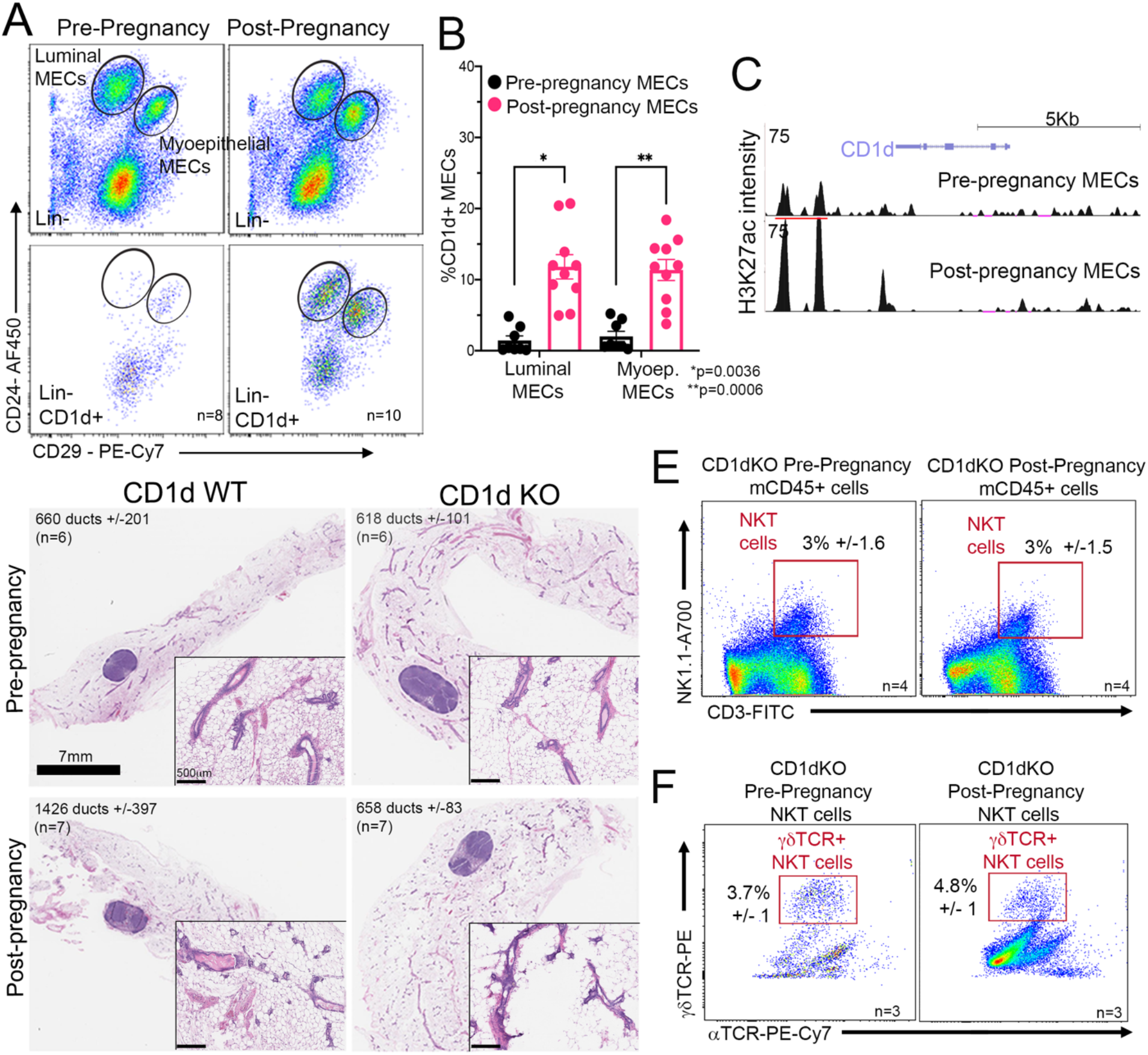
NKT expansion depends on CD1d expression on post-pregnancy MECs. (**A**) Flow cytometry analysis of myoepithelial and luminal MECs harvested from pre-pregnancy (and post-pregnancy mammary tissue, and their distribution based on CD1d cell-surface expression. (**B**) Flow cytometry quantification of CD1d+ MECs harvested from pre-pregnancy (black bars, n=8) and post-pregnancy (pink bars, n=10) mammary tissue. *p=0.0036 for luminal MECs and **p=0.0006 for myoepithelial MECs. (**C**) Genome browser tracks showing MACS-called, H3K27ac ChIP-seq peaks at the Cd1d genomic locus in FACS-isolated, pre- and post-pregnancy luminal MECs. (**D**) H&E stained histological images and duct quantification from mammary glands harvested from nulliparous (top left, n=6) and parous (bottom left, n=7) CD1d WT female mice, and nulliparous (top right, n=6) and parous (bottom right, n=7) CD1d KO female mice. p=0.86 for pre-pregnancy glands and p=0.78 for post-pregnancy glands. Scale: 7mm. Zoom in panels, scale 500μm. (**E**) Flow cytometry analysis of mammary resident CD45+ cells harvested from pre- and post-pregnancy CD1d KO female mice, and their distribution of NKT cells (NK1.1+CD3+). n=4 nulliparous and n=4 parous female mice. *p=0.3. (**F**) Flow cytometry analysis of α and γδ T-cell receptors (TCRs) of CD1d KO NKT cells from nulliparous (left, n=3) and parous (right, n=3) female mice. *p=0.5. For all analyses, error bars indicate standard error of mean across samples of the same experimental group. Statistically significant differences were considered with Student’s t-test *p*-value lower than 0.05 (p<0.05).

Gene expression analysis of scRNA-seq datasets and qPCR quantifications of FACS-isolated epithelial cells confirmed that post-pregnancy MECs express higher levels of *Cd1d* mRNA, supporting that pregnancy induced molecular alterations may represent the basis for the observed increase in the percentage of CD1d+ post-pregnancy MECs (**Fig. 1D and Supplementary Fig. S7D**). In agreement, we observed increase levels of the active transcription marker histone H3 lysine 27 acetylation (H3K27ac) at the Cd1d genomic locus in FACS-isolated post-pregnancy mammary MECs, suggesting that increased mRNA levels could be associated with parity-induced, epigenetic changes at the CD1d locus (**Fig. 3C**). These observations were confirmed in organoid systems that mimic the transcription and epigenetic alterations brought to MECs by pregnancy signals (Ciccone et al., 2020), where pregnancy hormones induced upregulation of Cd1d mRNA levels and increased H3K27ac levels at the CD1d locus (**Supplementary Fig. S7E-F**). Thus, pregnancy-associated signals may induce epigenetic alterations at the Cd1d gene locus, that subsequently associate with increased *Cd1d* mRNA and CD1d protein levels in post-pregnancy MECs.

To investigate whether CD1d expression is required for the expansion of NKT cells after parity, we analyzed mammary glands from CD1d KO mice, which bear reduced levels of activated NKT cells (Faunce et al., 2005; Macho-Fernandez and Brigl, 2015; Mantell et al., 2011). Mammary glands from nulliparous and parous CD1d KO mice displayed similar numbers of ductal structures and MEC populations as CD1d wild-type (WT) female mice, suggesting that loss of CD1d does not majorly alter mammary gland tissue homeostasis (**Fig. 3D**). Further flow cytometry analysis indicated no statistically significant changes in the percentage of NKT cells in mammary glands of nulliparous CD1d KO mice (2.2% +/− 0.8), compared to nulliparous CD1d WT mice (3% +/− 1.6) (**Fig.2A, left panel, and Fig.3E, left panel**). Conversely, we found a 7-fold decrease in the percentage of NKT cells in mammary tissue from fully involuted, parous CD1d KO female mice (3% +/− 1.5) compared to parous CD1d WT mammary tissue (26% +/− 4), supporting role of CD1d in regulating NKT activation (**Fig.2A, right panel, and Fig.3E, right panel**). Moreover, we found no difference in the abundance of NKT cells in glands from pre- and post-pregnancy CD1d KO female mice, consistent with lack of Cd1d expression reducing the activation of NKT cells (**Fig. 3E**). These results were supported by the analysis of an additional mice strain that is deficient in mature/activated NKT cells, due to the deletion of the histone-demethylase Kdm6 (Utx KO mouse model), which failed to detect an expansion of NKT cells post-pregnancy, supporting that pregnancy induces the expansion of mature/active subtypes of NKT cells (Beyaz et al., 2017) (**Supplementary Fig. S7G**). Moreover, NKT cells observed in post-pregnancy CD1d KO mammary tissue mainly expressed ββTCR on their surface, in contrast to the γδNKT cells observed in CD1d WT post-pregnancy glands, further confirming that loss of CD1d expression affects the expansion and activation of specific populations of NKT cells in post-pregnancy mammary tissue (**Fig. 3F**).

Collectively, our studies identify pregnancy-induced epigenetic changes that may control the expression of *Cd1d* mRNA in MECs, and elucidate a role for CD1d in mediating communication between the MECs and the immune cell population of γδTCR-expressing NKT cells, unique to post-pregnancy mammary glands.

### Lack of mammary oncogenesis is marked by NKT expansion and CD1d+ MECs in CAGMYC and Brca1 KO parous female mice

Parity resulted in the expansion of a specific population of γδNKT cells in the mammary gland in response to the up-regulation of CD1d on the surfaces of MECs, pointing to a mechanistic connection between pregnancy-associated MECs and immune cell biology. A pregnancy has also been demonstrated to induce molecular modifications to MECs associated with an oncogene-induced senescence response to cMYC overexpression, and thus suppression of MEC malignant transformation (Feigman et al., 2020). Therefore, we next investigated whether pregnancy-induced mammary cancer protection was associated with the expansion of NKT cells.

Flow cytometry analysis of pre- and post-pregnancy mammary tissue from cMYC overexpressing female mice (DOX-treated, CAGMYC model) demonstrated a 1.5-fold increase in the abundance of total CD3+ T-cells (**Supplementary Fig. S8A**). Increased levels of CD3+ T-cell expansion was also observed in mammary tissue transplanted with CAGMYC post-pregnancy MECs and organoid cultures derived from post-pregnancy CAGMYC MECs, both conditions previously demonstrated to lack mammary oncogenic development, and therefore suggesting a link between pregnancy-induced tumorigenesis inhibition and specific changes to the adaptive immune system (**Supplementary Fig. S8B-C**). This selective expansion of CD3+ T cells was further supported by the analysis of markers that define mammary resident neutrophils (Ly6G+), and mammary resident macrophages (CD206+), which were largely unchanged in mammary tissue transplanted with either pre- and post-pregnancy CAGMYC MECs (**Supplementary Fig. S8B**).

Further flow cytometry analysis identified a 6-fold increase in the percentage of NKT cells in mammary tissue from parous CAGMYC female mice, which predominantly expressed γδ TCRs (**Fig. 4A, Supplementary Fig. S8D**). No changes in the abundance of CD8+ T-cells or CD4+ T-cells was observed between mammary tissue from nulliparous and parous CAGMYC female mice, supporting the parity-induced expansion of γδNKT cells (**Supplementary Fig. S8E-F**), and suggesting that specific constituents of the mammary immune microenvironment may control mammary tumorigenesis. In agreement, we also found a 5-fold higher percentage CD1d+ luminal MECs in post-pregnancy mammary tissue, thus linking gain of CD1d expression and the expansion of γδTCR-expressing NKT cells, which may collectively play a role in blocking tumorigenesis (**Fig. 4B**).

**Figure 4.**
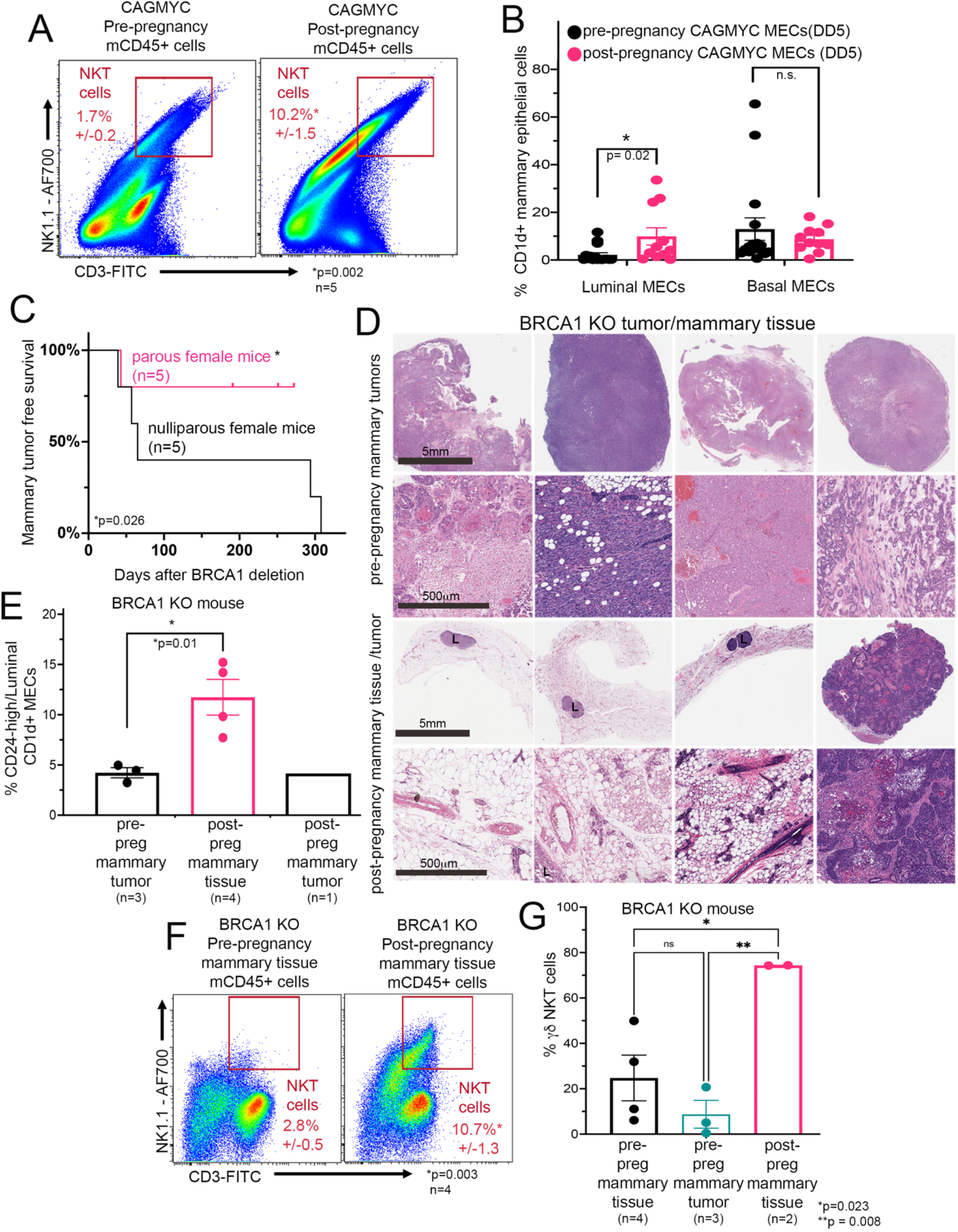
Lack of mammary oncogenesis is marked by NKT expansion and CD1d+ MECs in CAGMYC and Brca1 KO parous female mice. (**A**) Flow cytometry analysis of mammary resident NKT cells (CD45+NK1.1+CD3+) from DOX-treated nulliparous (left panel, n=5) and parous (right panel, n=5) CAGMYC female mice. *p=0.002. (**B**) Flow cytometry quantification of CD1d+ luminal and myoepithelial MECs from DOX-treated nulliparous (left panel, n=16) and parous (right panel, n=11) CAGMYC female mice. *p=0.02. (**C**) Mammary tumor-free survival plot of nulliparous (black line, n=5) and parous (pink line, n=5) Brca1 KO female mice. (**D**) H&E stained histological images from mammary tissue and mammary tumor harvested from nulliparous (top panels) and parous (bottom panels) Brca1 KO female mice. (**E**) Flow cytometry quantification of CD1d+ CD24^high^ luminal MECs from Brca1 KO pre-pregnancy mammary tumors (black bar, n=3), Brca1 KO post-pregnancy healthy mammary tissue (pink bar, n=4), and Brca1 KO post-pregnancy mammary tumor (blue bar, n=1). *p=0.02. (**F**) Flow cytometry analysis of mammary resident NKT cells in normal mammary tissue from nulliparous, tumor-bearing, Brca1 KO female mice (left panel, n=4) and normal mammary tissue from healthy parous Brca1 KO female mice (right panel, n=4). *p=0.003. (**G**) Quantification of γδNKT cells in normal mammary tissue from nulliparous, tumor-bearing, Brca1 KO female mice (black bar panel, n=4), in mammary tumor tissue from nulliparous Brca1 KO female mice (blue bar, n=3), and in normal mammary tissue from healthy parous Brca1 KO female mice (black bar panel, n=2). *p=0.023 and **p=0.008. For all analyses, error bars indicate standard error of mean across samples of the same experimental group. Statistically significant differences were considered with Student’s t-test *p*-value lower than 0.05 (p<0.05).

cMYC overexpression is present in approximately 60% of basal-like breast cancers, with cMYC gain of function commonly found in BRCA1 mutated breast cancers (Chen and Olopade, 2008; Grushko et al., 2004). Interestingly, women harboring *BRCA1* mutations with a full-term pregnancy before the age of 25 benefit from pregnancy-induced breast cancer protection (Medina et al., 2004; Terry et al., 2018). Therefore, we developed an inducible mouse model of *Brca1* loss of function, for the purpose of investigating how pregnancy-induced changes influences Brca1 null mammary tumor development. In this model, tamoxifen (TAM) induces homozygous loss of Brca1 function in cells that express the cytokeratin 5 gene (KRT5+ cells), which include MECs (dos Santos et al., 2013), cells from gastrointestinal tract (Sulahian et al., 2015), reproductive organs (Ricciardelli et al., 2017), and additional epithelial tissue (Castillo-Martin et al., 2010; Majumdar et al., 2012), in p53 heterozygous background (*Krt5*^CRE-ERT2^*Brca1*^fl/fl^*p53*^−/+^, hereafter referred as Brca1 KO mouse).

Nulliparous Brca1 KO mice exhibited signs of mammary hyperplasia approximately 12 weeks post TAM treatment, which gradually progressed into mammary tumors at around 20 weeks after Brca1 deletion (**Supplementary Fig.S9A-B**). Brca1 KO mammary tumors display cellular and molecular features similar to those previously described in human breast tissue from BRCA1 mutant carriers and animal models of Brca1 loss of function, including high EGFR and KRT17 protein levels and altered copy number variation marked by gains and losses of genomic regions (**Annunziato et al. Nat Comm. 2019**) (**Supplementary Fig.S9C-D**).

To investigate the effects of pregnancy on the mammary immune microenvironment and mammary oncogenesis, age matched, TAM-treated, Brca1 KO nulliparous female mice, and parous Brca1 KO female mice (1 pregnancy, 21-days of gestation, 21-days of lactation/nursing, and 40-days post offspring weaning) were monitored for tumor development (**Supplementary Fig.S10A**). Our study demonstrated that 100% of all nulliparous Brca1 KO female mice (5 out of 5 mice) developed mammary tumors, compared to only 20% of the parous Brca1 KO female mice that developed mammary tumors (1 out of 5), thus indicating that a full pregnancy cycle decreases the frequency of Brca1 KO mammary tumors by 80% (**Fig. 4C-D**).

Histo-pathological analysis suggested that pre-pregnancy mammary tumors were quite diverse as previously reported for tumors from Brca1 KO mice (Brodie et al., 2001). These included poorly differentiated tumors, such as micro-lobular carcinomas with squamous trans-differentiation (**Fig. 4D – top rows, far left panel**), medullary like carcinomas (**Fig. 4D – top rows, right panel**), and solid carcinomas resembling high-grade invasive ductal carcinoma (IDC) in humans (**Fig. 4D – top rows, left and far right panels**). Accordingly, the only tumor-bearing parous BRCA1 KO female mouse developed a poorly differentiated carcinoma with extensive squamous trans-differentiation and with extensive necrosis, also previously reported for tumors from Brca1 KO mice (**Fig. 4D – bottom rows, far right panels**). Additional histo-pathological analysis confirmed that mammary tissues from the remaining parous Brca1 KO female mice (4 out of 5) were largely normal (**Fig. 4D – bottom rows, far left, left and right panels and Supplementary Fig. S10B**). Immunofluorescence analysis confirmed that both pre-pregnancy mammary tumors and post-pregnancy normal mammary tissue were indeed deficient for KRT5+BRCA1+ epithelial cells, indicating that the lack of mammary tumors in parous female mice was not due to inefficient Brca1 deletion (**Supplementary Fig. S11A**).

Flow cytometry analysis of Brca1 KO MECs demonstrated a progressive loss of myoepithelial cells in tumor tissue from nulliparous (2.5-fold) and parous (2-fold) Brca1 KO female mice, defined by an increase in the percentage of CD24^high^CD29^low^ luminal-like MECs, (**Supplementary Fig. S11B**). These results suggest that tumor progression in this model is accompanied by changes to the population of CD24^high^ MECs, which has been associated with poor clinical outcomes in patients with triple negative breast cancer (Chan et al., 2019). Further cellular analysis indicated a 2.7-fold increased on the percentage of CD24^high^/luminal cells CD1d+ cells in healthy, post-pregnancy Brca1 KO mammary tissue compared to tissue from tumor-bearing nulliparous Brca1 KO mice and parous Brca1 KO mice, supporting that parity induces the expression of CD1d at the surface of MECs (**Fig.4E**).

Given the increased levels of CD1d expression at the surface of post-pregnancy Brca1 KO MECs, we next investigated the presence of NKT cells in mammary tissue from nulliparous and parous Brca1 KO female mice. Flow cytometry analysis demonstrated a 3.8-fold increase in the percentage of NKT cells in healthy, post-pregnancy Brca1 KO mammary tissue compared to non-affected normal mammary tissue from tumor-bearing, nulliparous Brca1 KO mice and parous Brca1 KO mice (**Fig.4F and Supplementary Fig. S11C**). Additional flow cytometry analysis demonstrated that approximately 70% of total NKT cells from healthy, post-pregnancy Brca1 KO mammary tissue expressed γδTCR, in marked contrast to NKT cells from healthy (2.7%) and tumor mammary tissue (8.6%) from nulliparous Brca1 KO mice (**Fig.4G**).

Collectively, our findings show that pregnancy-induced gain of CD1d expression at the surface of MECs and expansion of NKT cells associates with lack of mammary oncogenesis in response to cMYC overexpression or loss of Brca1 function, thus supporting to the link between pregnancy-induced molecular changes, mammary tissue immune alteration, and inhibition of mammary tumorigenesis in clinically relevant mouse models.

### Functionally active NKT cells are required to block malignant progression of post-pregnancy MECs

Given that we demonstrated that pregnancy-induced changes block mammary oncogenesis in two distinct models (**Fig.4**), and that cMYC gain of function is commonly found in BRCA1 mutated breast cancers, we utilized the cMYC overexpression model to further characterize the effects of the immune microenvironment on the malignant progression of post-pregnancy MECs. Analysis of fat-pad transplantations into severely immune deficient NOD/SCID female mice, which lack T-cells, B-cells, NK and NKT cells, indicated that 100% of mammary tissue injected with pre-pregnancy (n=5) or post-pregnancy (n=5) CAGMYC MECs developed adeno-squamous-like carcinomas with acellular lamellar keratin, high levels of cell proliferation (Ki67 staining), and increased collagen deposition (Trichrome blue staining) (**Supplementary Fig. S12A-C**). Therefore, NKT cells, or associated adaptive immune cells, are required for the parity associated protection from oncogenesis in the CAGMYC model.

Bulk RNA-seq analysis demonstrated that post-pregnancy CAGMYC MECs transplanted into the fat-pad of NOD/SCID female mice were less effective at activating the expression of canonical cMYC targets and estrogen response genes, compared to transplanted pre-pregnancy CAGMYC MECs, in agreement with the previously reported transcriptional state of post-pregnancy CAGMYC MECs (Feigman et al., 2020) (**Supplementary Fig. S12D**). We also found that organoid cultures derived from post-pregnancy CAGMYC MECs transplanted into NOD/SCID female mice retained a senescent-like state, characterized by reduced p300 protein levels and moderately increased p53 protein levels, in agreement with the previously reported senescence state of post-pregnancy CAGMYC MECs (Feigman et al., 2020) (**Supplementary Fig. S12E**). Together, these findings indicate that oncogenic progression of post-pregnancy CAGMYC MECs is associated with the immune deficient mammary microenvironment of NOD/SCID mice.

While our investigation of post-pregnancy CAGMYC MECs that were transplanted into the mammary tissue of immunosuppressed animals alluded to the importance of a robust immune system in blocking mammary tumor development, it did not uncouple whether functionally active NKT cells, or CD1d expression at the surface of MECs, act to block oncogenesis in post-pregnancy mammary tissue. Therefore, to determine whether signaling between CD1d+ MECs and NKT cells is critical for the development of mammary oncogenesis after pregnancy, we developed a double transgenic mouse model, by crossing the DOX-inducible CAGMYC mice into a CD1d KO background, hereafter referred as CAGMYC CD1d KO.

Tissue histology analysis indicated that mammary tissue from DOX-treated, nulliparous and parous CAGMYC CD1d KO female mice showed signs of tissue hyperplasia with atypia and abnormal ductal structures, demonstrating that loss of Cd1d expression is accompanied by mammary oncogenesis in a parity-independent fashion (**Fig. 5A, left and far right panels and Supplementary Fig. S13A**). Conversely, analysis of DOX-treated, CAGMYC CD1d WT mice showed that mammary tissue from parous female mice lacked malignant lesions in response to cMYC overexpression (**Fig. 5A, right panels and Supplementary Fig. S13A**). Flow cytometry analysis showed a lack of NKT cells in mammary tissue from both nulliparous and parous CAGMYC CD1d KO female mice, in marked contrast to the observed expansion of γδ NKT cells in healthy post-pregnancy CAGMYC CD1d WT mammary glands that lacked tissue hyperplasia development, suggesting that CD1d expression may control pregnancy-induced expansion/activation of NKTs, and thus block of mammary tumorigenesis. (**Supplementary Fig. S13B and Fig.4A**). To further determine whether loss of CD1d expression underlies the malignant transformation of post-pregnancy CAGMYC MECs, we performed mammary transplantation assays of CAGMYC CD1d KO MECs into the fat-pad of syngeneic animals (CD1d WT female mice). We found that 100% of mammary tissue injected with pre-pregnancy CAGMYC CD1d KO MECs and 70% of mammary glands injected with post-pregnancy CAGMYC CD1d KO MECs developed signs of malignant lesions, supporting that the loss of CD1d expression impacts pregnancy-induced breast cancer protection (**Fig. 5B - black font**, and **Supplementary Fig. S13C-D**). This last observation was in marked contrast to the finding in glands injected with post-pregnancy CAGMYC CD1d WT MECs, which as previously reported did not present signs of malignant transformation (Feigman et al., 2020) (**Fig. 5B, blue font and Supplementary Fig. S13E-F**).

**Figure 5.**
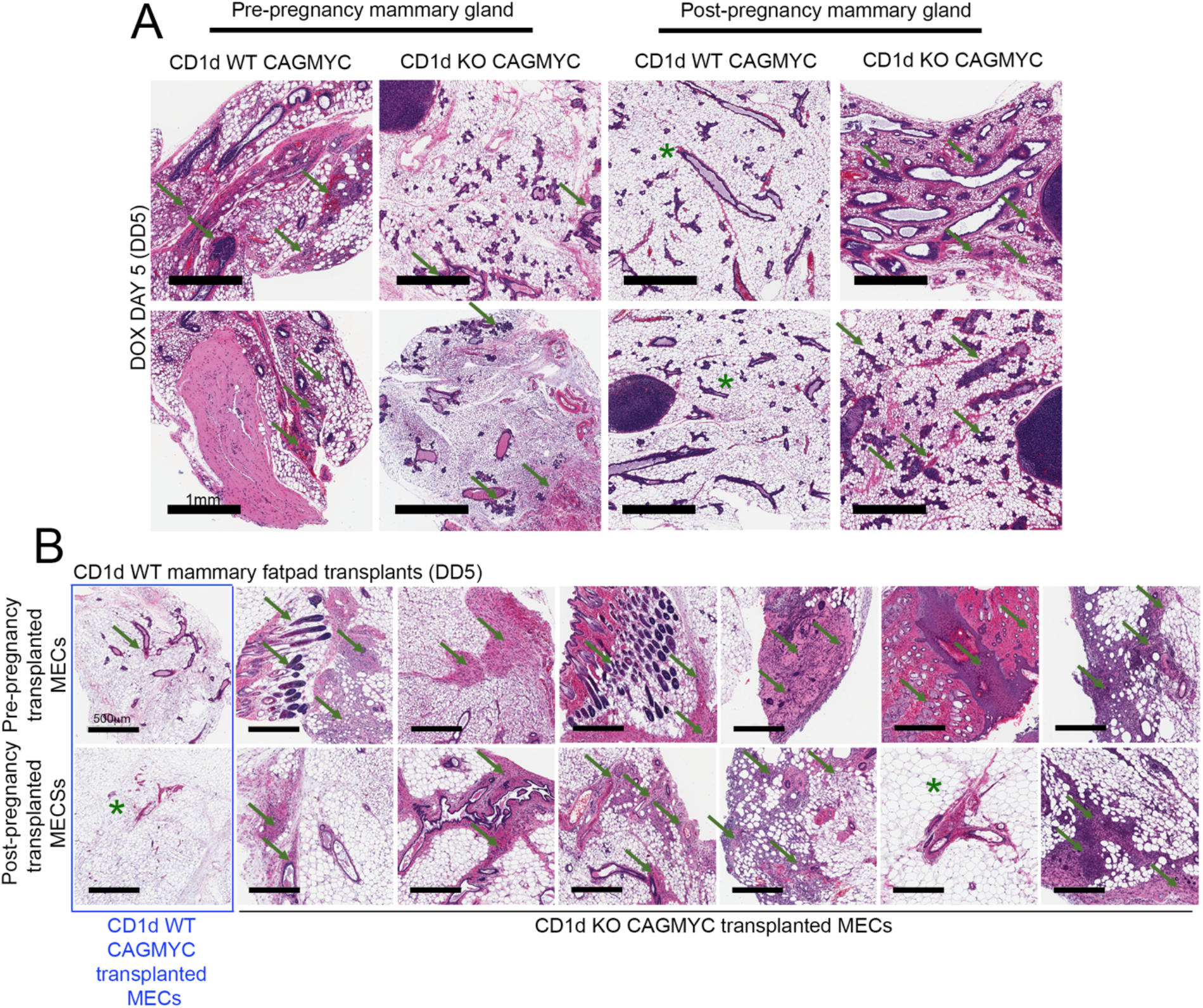
Functionally active NKT cells are required to block malignant progression of post-pregnancy MECs. (A) H&E stained histological images of mammary tissue harvested from DOX-treated (DD5), nulliparous CD1d WT CAGMYC (far left panels), nulliparous CD1d KO CAGMYC (left panels), parous CD1d WT CAGMYC (right panels), and parous CD1d KO CAGMYC (far right panels) female mice. Green arrows indicate signs of malignant lesions/mammary hyperplasia. Green asterisks indicate normal-like ductal structures. (B) H&E stained histological images of DOX-treated, CD1d WT mammary tissue transplanted with pre-pregnancy CD1d WT CAGMYC MECs (blue font, top far left panel), pre-pregnancy CD1d KO CAGMYC MECs (black font, top panel), post-pregnancy CD1d WT CAGMYC MECs (blue font, bottom far left panel), or post-pregnancy CD1d KO CAGMYC MECs (black font, bottom panel). Green arrows indicate signs of malignant lesions/mammary hyperplasia. Green asterisks indicate normal-like ductal structures.

Altogether, these results suggest that loss of CD1d, with concomitant loss of pregnancy-induced expansion of NKT cells, supports the development of mammary malignant lesions, independently of parity. Moreover, our study elucidates that parity blocks the malignant transformation of MECs, both by inducing cell-autonomous, epigenetic alterations within the MECs, and non-autonomous, communication between CD1d+ MECs cells and NKT cells in the mammary gland.

## Discussion

In mammals, reprogramming of the immune system is initiated after birth, and continues throughout the lifespan of an individual due to exposure to pathogens, hormonal fluctuations, and aging. This dynamic reprogramming is part of an immune surveillance system that detects abnormal cells across many tissues, helping to prevent cancer. Here, we characterized a population of mammary resident NKT immune cells in post-pregnancy mammary tissue, and it’s role on inhibiting mammary oncogenesis

Our findings suggest that post-pregnancy mammary homeostasis does not rely on the presence of γδNKT cells, given the largely normal histology and cellular content of mammary tissue in mice deficient for this cell type. It is possible that NKT cells expand in response to the re-setting of whole-body immunity post-partum, with the child-bearing event providing signals that alters antigens across all maternal tissues as well as expanding specific immune cell populations. γδNKT cells have been found in the pregnant uterus across many mammalian species, linking NKT specialization and the pregnancy cycle (Mincheva-Nilsson, 2003). Our results support that the expansion of NKT cells was predominantly observed in post-lactating, post-involution tissue, thus suggesting that the immune reprogramming of mammary tissue takes place after giving birth. In addition to the NKT cell population expansion, parity also promotes a modification of the TCR repertoire in NKT cells. γδT-cells reside within the normal breast, and their presence has been associated with a better prognosis during triple-negative breast cancer development (Wu et al., 2019). Here we report that pregnancy-induced changes in TCR expression was specific to NKT cells, given that we did not find pregnancy-induced TCR rearrangements in CD8+NK1.1-immune cells, pointing to the specific engagement of NKT-lineages during pregnancy-induced mammary development.

Several other immune subtypes have been described to be enriched in mammary tissue during gestation, lactation and post-pregnancy involution stages of mammary gland development. These studies identified alterations in leukocyte interaction with mammary ductal structures, as well to specific transcriptional changes, suggesting that cell interaction and cellular identity of mammary resident cells are affected by pregnancy-induced development (Dawson et al., 2020; Hitchcock et al., 2020). Our analysis of leukocytes, specifically macrophages and neutrophils, did not show alterations in cell abundance, neither in mammary tissue from healthy parous female mice, nor in post-pregnant CAGMYC mammary tissue lacking malignant lesions. Moreover, we found that CD1d expression on the surface of total CD45+ mammary resident immune cells were not altered by parity, thus supporting a role for post-pregnancy CD1d+ MECs in regulating CD1d-dependent NKT cells. However, given that leukocytes have been implicated in the activation of NKT cells (Macho-Fernandez and Brigl, 2015; Rizvi et al., 2015), it is possible that molecular alterations, rather than changes to cellular abundance or antigen presentation, could play a role in inducing or sustaining the population of NKT cells in post-pregnancy mammary tissue.

Our studies also provide evidence linking pregnancy-induced immune changes with the inhibition of mammary oncogenesis. Our previous research focused on how post-pregnancy MECs assume a senescence-like state in response to cMYC overexpression, an oncogene-induced response that activates the immune system via the expression of senescence-associated genes (Braig and Schmitt, 2006). Here, we found that CD1d expression at the surface of post-pregnancy MECs, and the presence of γδNKT cells were linked with the inhibition of mammary oncogenesis in two independent models of breast cancer, illustrating how epithelial and immune cells communicate to support pregnancy-induced mammary cancer prevention. Given that NKT cells were previously shown to interact with senescent cells, it is possible that pregnancy-induced activation of CD1d expression and NKT cell expansion represent additional responses to oncogene-induced cellular senescence (Kale et al., 2020).

Women completing a full-term pregnancy before the age of 25 have a substantially reduced breast cancer risk, by approximately one-third (Medina et al., 2004). This benefit applies to the risk of all breast cancer subtypes, including those from women harboring *BRCA1* mutations (Terry et al., 2018). Thus, our findings supporting a role for pregnancy in inhibiting the development of Brca1 KO mammary tumors lends a clinical relevance to our studies. Interestingly, mammary tumor from parous Brca1 KO female mouse was associated with low abundance of γδNKT cells and CD1d+ MECs, suggesting that loss of the pregnancy-induced epithelial to immune microenvironment communication may support mammary tumorigenesis. In agreement, the genetically engineered loss of CD1d expression, with a consequent deficiency in activated NKTs, supported the malignant progression of cMYC overexpressing MECs, thus further illustrating a link between epithelial and immune cells in supporting pregnancy-induced mammary cancer prevention.

Our findings are based on studies performed in mice that became pregnant at a young age (~8 weeks old), which reinforced pregnancy-induced changes to epithelial cells, and their effect on immune recruitment and oncogenesis inhibition. However, it remains unclear why such strong, pregnancy-induced changes do not fully prevent the development of breast cancer (Nichols et al., 2019). It has been suggested that specific mammary epithelial clones with oncogenic properties reside within the mammary tissue after pregnancy, and may give rise to late-onset mammary oncogenesis in aged mice (Li et al., 2020b). It is possible that such populations of rare MECs lose some of their pregnancy-induced molecular signatures over time, thereby bypassing oncogene-induced senescence and immune recognition, and ultimately developing into mammary tumors. Moreover, and given that pregnancy-induced breast cancer protection becomes apparent ~5-8-years after pregnancy, it is possible that additional immune reprogramming induced by genetic makeup, age at pregnancy, and/or overall post-partum health, may further modify breast tissue and erase pregnancy-induced changes that inhibit breast cancer development.

Nonetheless, the connection between pregnancy, immunity, and oncogenesis could be used to develop therapies to block cancer development. For example, strategies could be developed to induce NKT expansion in the absence of a true pregnancy. Indeed, a series of preclinical models have been developed to optimize the delivery of CD1d stimulatory factors, such as aGalcer and KRN7000, and induce expansion of NKT cells (Zhang et al., 2019). Such strategies are mostly side-effect free, and could be used in cases of high cancer risk, including in the event of genetic alterations that affect Brca1 function and/or family history of breast cancer. Additionally, the characterization of specific, pregnancy-induced TCR rearrangements could be leveraged in CAR-NKT immunotherapy, for example, which could also efficiently target disease that has already developed. Collectively, such strategies could also improve breast health, nursing experience, and decrease cancer risk in women that experience their first pregnancy after 35 years of age, when they are at greater risk of requiring medical intervention to improve milk production and breastfeeding assistance and to develop breast cancer.

## Acknowledgements

This work was performed with assistance from CSHL Animal Facility, the CSHL Tissue Histology Shared Resources, the CSHL NextGen Sequencing Shared Resources, the CSHL Single Cell Shared Resources, the CSHL Flow Cytometry Shared Resources, and the CSHL Microscopy Shared Resources, which are supported by the CSHL Cancer Center Support Grant 5P30CA045508. This work was financially supported by the CSHL and Northwell Health affiliation, the CSHL and Simons Foundation Award, the Rita Allen Scholar Award, the Pershing Square Sohn Prize for Cancer Research, the Breast Cancer Research Foundation, the NIH/NCI grant R01CA248158-01, and the NIH/NIA grant R01 AG069727-01 (C.O.D.S.). Whole genome sequencing (CNV analysis) was performed with financial support provided to Dr. Michael Wigler by The Breast Cancer Research Foundation (BCRF-19-174) and Simons Foundation, Life Sciences Founder Directed Giving-Research (519054). We would like to thank Dr. Douglas Fearon for critical feedback, and Mrs. Shih-Ting Yang for the mouse illustrations.

## Author contributions

C.O.D.S. designed and supervised the research; C.O.D.S, A.V.H.S., M.A.M., M.J.F., and S.L.C. wrote the manuscript. A.V.H.S., M.A.M., M.J.F., C.C., S.L.C., M.F.C., M.C.T, and M.V. performed experiments and analyzed results. M.A.M, and M.C.T. performed bioinformatics analyses. S.L., and J.K., performed and analyzed whole genome sequencing (CNV analysis). S.B. provided reagents and critical feedback. J.E.W. performed histopathological analysis.

## Resource availability

### Materials Availability

All unique/stable reagents generated in this study are available from the Lead Contact with a completed Materials Transfer Agreement.

### Data and Code Availability

scRNA-seq, RNA-seq, ATAC-seq datasets were deposited into BioProject database under number PRJNA708263, and will be made available upon manuscript acceptance/publication. Whole genome sequencing datasets were deposited under number SUB10186897. Results shown in Figure 1 (pre-pregnancy scRNA-seq) were previously deposited under number SUB8429356 (data pending release). Results shown in Supplementary Fig. S2C (pre- and post-pregnancy RNA-seq), Fig.3C (pre- and post-pregnancy H3K27ac ChIP-seq) were previously deposited in the BioProject database under numbers PRJNA192515 [https://www.ncbi.nlm.nih.gov/bioproject/?term=PRJNA192515] and PRJNA544746 [ https://www.ncbi.nlm.nih.gov/bioproject/PRJNA544746]. Results shown on Supplementary Fig. S7F (H3K27ac Cut&Run of organoid cultures) was previously deposited in the BioProject database under number PRJNA656955 (https://www.ncbi.nlm.nih.gov/sra/?term=PRJNA656955).

## Experimental model and subjects details

### Animal Studies

All experiments were performed in agreement with approved CSHL Institutional Animal Care and Use Committee (IACUC). All animals were housed at a 12 hour light/12 hour dark cycle, with a controlled temperature of 72°F and 40-60% of humidity. Balb/C female mice were purchased from The Jackson Laboratory and Charles River. RAG1 KO mice (B6.129S7-Rag1^tm1Mom/^J, IMSR Cat# JAX:002216, RRID:IMSR_JAX:002216) were purchased from The Jackson Laboratory. VavCre UTX KO were generated as previously described (Beyaz et al., 2017). CXCR6-KO-EGFP-KI mice (B6.129P2-Cxcr6^tm1Litt^/J, IMSR Cat# JAX:005693, RRID:IMSR_JAX:005693) were purchased from The Jackson Laboratory. CAGMYC transgenic mouse strain was generated as previously described (Feigman et al., 2020). CD1d KO CAGMYC transgenic mouse stain was generated by crossing CD1d KO (C.129S2-Cd1^tm1Gru^/J, IMSR Cat# JAX:003814, RRID:IMSR_JAX:003814) mice with CAGMYC mice. Krt5^CRE-ERT2^Brca1^fl/fl^p53^het^(Brca1 KO) transgenic mouse strain was generated by crossing Blg^CRE^Brca1^fl/fl^p53^het^ transgenic mouse strain (Trp53^tm1Brd^Brca1^tmAash^Tg(B-cre)74Acl/J, IMSR Cat# JAX:012620, RRID:IMSR_JAX:012620) with Krt5^CRE-ERT2^ transgenic mouse strain (B6N.129S6(Cg)-Krt5^tm1.1(cre/ERT2)Blh^/J, IMSR Cat# JAX:029155, RRID:IMSR_JAX:029155).

### Methods details

Full description of methods is provided as Supplementary Information.

