## Supplemental Methods for "Parity-induced changes to mammary epithelial cells control NKT cell expansion and mammary oncogenesis"

**Antibodies.** All antibodies were purchased from companies as indicated bellow and used without further purification. Antibodies for lineage depletion: biotinylated anti-CD45 (Thermo Fisher Scientific Cat# 13-0451-85, RRID:AB_466447), biotinylated anti-CD31 (Thermo Fisher Scientific Cat# 13-0311-85, RRID:AB_466421), biotinylated anti-Ter119 (Thermo Fisher Scientific Cat# 13-5921-85, RRID:AB_466798) and biotinylated anti-CD34 (Thermo Fisher Scientific Cat# 13-0341-82, RRID:AB_466425). Antibodies for cell surface flow cytometry: eFluor 450 conjugated anti-CD24 (Thermo Fisher Scientific Cat# 48-0242-82, RRID:AB_1311169), PE-Cy7 conjugated anti-CD29 (BioLegend Cat# 102222, RRID:AB_528790), 7-AAD viability staining solution (BioLegend, #420404, RRID:SCR_020993), PerCP-Cy5.5 conjugated anti-CD1d (BioLegend Cat# 123514, RRID:AB_2073523), PE conjugated anti-CD1d (BioLegend Cat# 140805, RRID:AB_10643277), APC conjugated anti-CD45 (BioLegend Cat# 103112, RRID:AB_312977), FITC conjugated anti-CD3 (BioLegend Cat# 100204, RRID:AB_312661), Alexa Fluor 700 conjugated. anti-NK1.1 (BioLegend Cat# 108730, RRID:AB_2291262), APC/Cy7 conjugated anti-CD8 (BioLegend Cat# 100714, RRID:AB_312753), PE conjugated anti-TCR $\gamma/\delta$ (BioLegend Cat# 118108, RRID:AB_313832), APC conjugated anti-TCR $\beta$ (BioLegend Cat# 109212, RRID:AB_313435), APC conjugated anti-H-2Kb (BioLegend Cat# 116517, RRID:AB_10568693), Pacific Blue conjugated anti-I-Ab (BioLegend Cat# 116421, RRID:AB_10613291), Brilliant Violet 421 conjugated anti-CD206 (BioLegend Cat# 141717, RRID:AB_2562232), Alexa Fluor 700 conjugated anti-Ly6G (BioLegend Cat# 127621, RRID:AB_10640452). Antibodies for intracellular flow cytometry: PE conjugated anti-IFN$\gamma$ (BioLegend Cat# 505808, RRID:AB_315402), Pacific Blue conjugated anti-T-bet (BioLegend Cat# 644807, RRID:AB_1595586). Antibodies for negative controls: eFluor 450 conjugated mouse IgG (Thermo Fisher Scientific Cat# 48-4015-82, RRID:AB_2574060), FITC conjugated rat IgG (Thermo Fisher Scientific Cat# 11-4811-85, RRID:AB_465229), and PE-Cy7 conjugated mouse IgG (BioLegend Cat# 405315, RRID:AB_10662421). Antibody for MaSC enrichment: biotinylated anti-CD1d (BioLegend Cat# 123505, RRID:AB_1236543). Antibodies for Western Blot: anti-p300 antibody (Santa Cruz Biotechnology Cat# SC-585, RRID:AB_2231120), anti-Vinculin antibody (Abcam Cat# ab129002, RRID:AB_11144129), anti-p53 antibody (Leica Biosystems Cat# P53-CM5P, RRID:AB_2744683), goat anti-rabbit IgG HRP (Abcam Cat# ab6721, RRID:AB_955447) and goat anti-mouse IgG HRP (Abcam Cat# ab97051, RRID:AB_10679369). Antibodies for Immunohistochemistry (IHC) staining: anti-Cytokeratin 5 (KRT5) (BioLegend Cat# 905501, RRID:AB_2565050), anti-Cytokeratin 7/17 (KRT7/17) (Santa Cruz Biotechnology Cat# sc-8421, RRID:AB_627856), anti-EGFR (Santa Cruz Biotechnology Cat# sc-373746, RRID:AB_10920395), anti-AR (Santa Cruz Biotechnology Cat# sc-7305, RRID:AB_626671), and anti-Ki67 (Spring Bioscience Cat# M3062, RRID:AB_11219741). Antibodies for Immunofluorescence (IF) staining: Alexa Fluor 647 conjugated anti-Cytokeratin 5 (KRT5) (Abcam Cat# AB193895, RRID:AB_2728796), unconjugated rabbit anti-BRCA1 (Bioss Cat# bs-0803R, RRID:AB_10858843), Alexa Fluor 568 conjugated goat anti-rabbit IgG (Thermo Fisher Scientific Cat# A-11036, RRID:AB_10563566), Alexa Fluor 488 conjugated anti-GFP (BioLegend Cat# 338007, RRID:AB_2563287), Alexa Fluor 405 conjugated anti-Cytokeratin 8 (KRT8) (Abcam Cat# ab210139, RRID:AB_2890924).

**Mammary gland isolation.** Female mice classified as Pre-pregnancy (nulliparous, never pregnant), Post-pregnancy (parous, 1 gestation cycle, 21 days of lactation and 40 days of involution post offspring weaning), were housed together for 1-2 weeks to allow for estrous cycle synchronization prior to mammary gland isolation. For the experiments utilizing exposure to pregnany hormones (EPH), never pregnant female mice (~8 weeks old) were were implanted with 21 days-slow-release estrogen and progesterone pellets (17$\beta$-Estradiol (0.5 mg/pellet) + Progesterone (10 mg/pellet) – Innovative Research of America Cat# HH-112) prior to mammary gland isolation (at D12 post pellet implantation). Females classified as involution D15 had 1 gestation cycle, 21 days of lactation and 15 days of involution post offspring weaning. In all cases, mammary gland isolation was performed as previously described (dos Santos et al., 2013). In short, mammary glands (one to four pairs per mouse) were harvested, minced, and incubated for 2 hours with 1x Collagenase/Hyaluronidase (10x solution, Stem Cell Technology #07912) in RPMI 1640 GlutaMAX supplemented with 5% FBS. Digested mammary gland fragments were washed with cold HBSS (Thermo Fisher Scientific #14175103) supplemented with 5% FBS, followed by incubation with TrypLE Express (Thermo Fisher Scientific #12604-013) and an additional HBSS wash. Cells were incubated with 2 mL of Dispase (Stem Cell Technology #07913) supplemented with 40 µL DNAse I (Sigma #D4263) for 2 minutes and then filtered through a 100µm Cell Strainer (BD Falcon #352360). The single cell suspension was incubated with lineage depletion antibodies and loaded onto a MACS magnetic column (Miltenyi Biotec #130-042-401). Lineage negative, flow-through cells (epithelial cells) were utilized for flow cytometry, and transcriptomic analysis. Lineage positive cells (immune cells) were eluted from column with 3ml of MACS buffer and utilized for flow cytometry, transcriptomic and epigenomic analysis. For cell analysis, Dual Fortessa II cell analyzer (BD Biosciences) was used. Data analysis was performed using BD FACSDiva Software (RRID:SCR_001456) or FlowJo (FlowJo, RRID:SCR_008520). Statistically significant differences were considered with Student's t-test *p*-value lower than 0.05 ($p<0.05$).

**Flow cytometry gating analysis.** Mammary resident cells (epithelial and non-epithelial) were harvested from both top and bottom mammary glands, and analyzed according to the bellow indicated strategy. For all flow cytometry analysis an average of 300,000 cells live cells (7-AAD negative) were recorded.

a) General innate immune cells analysis strategy:

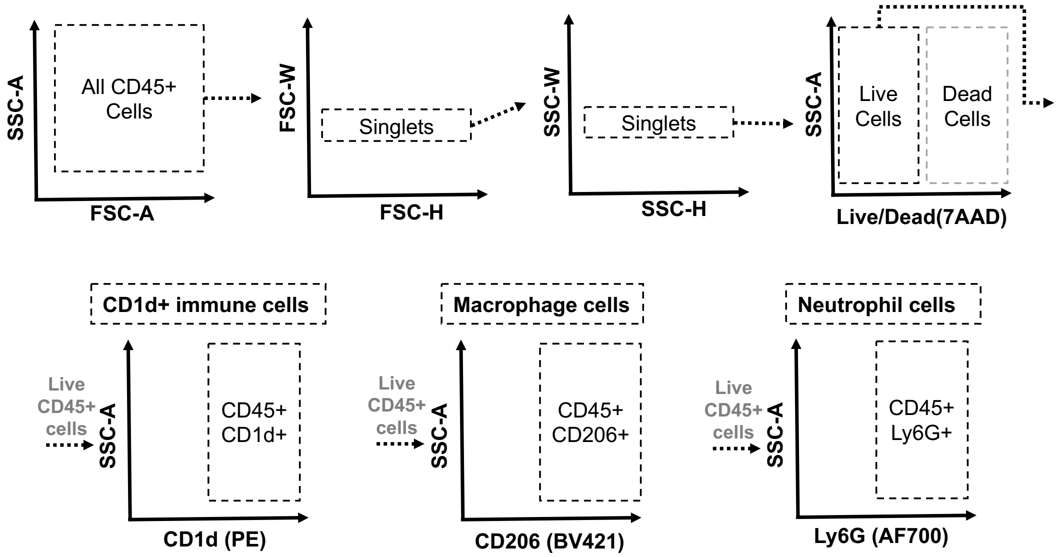

b) General adaptive immune cells analysis strategy:

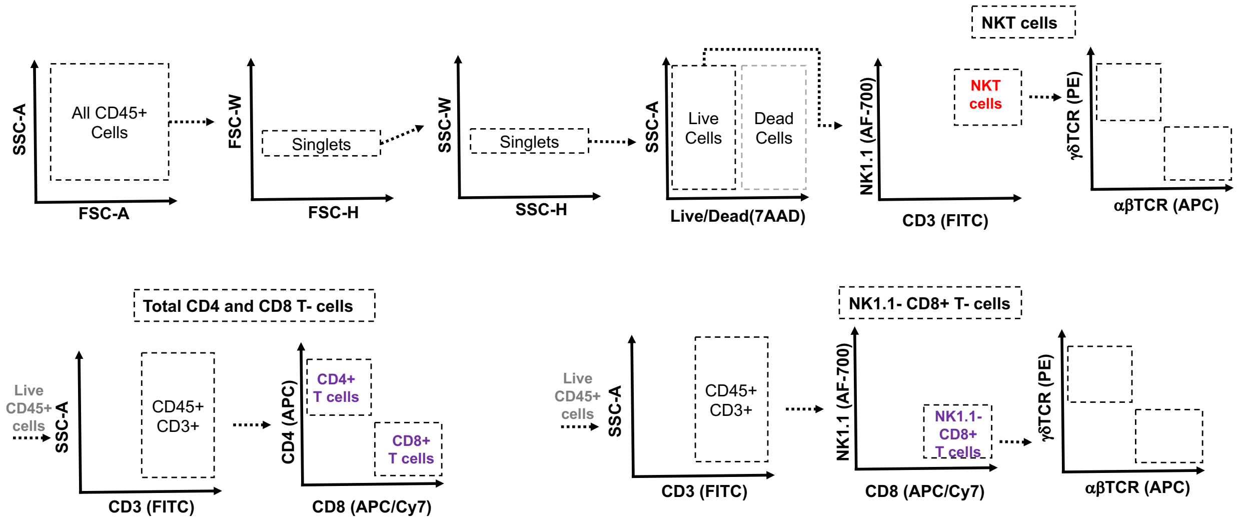


c) NKT intracellular analysis strategy:

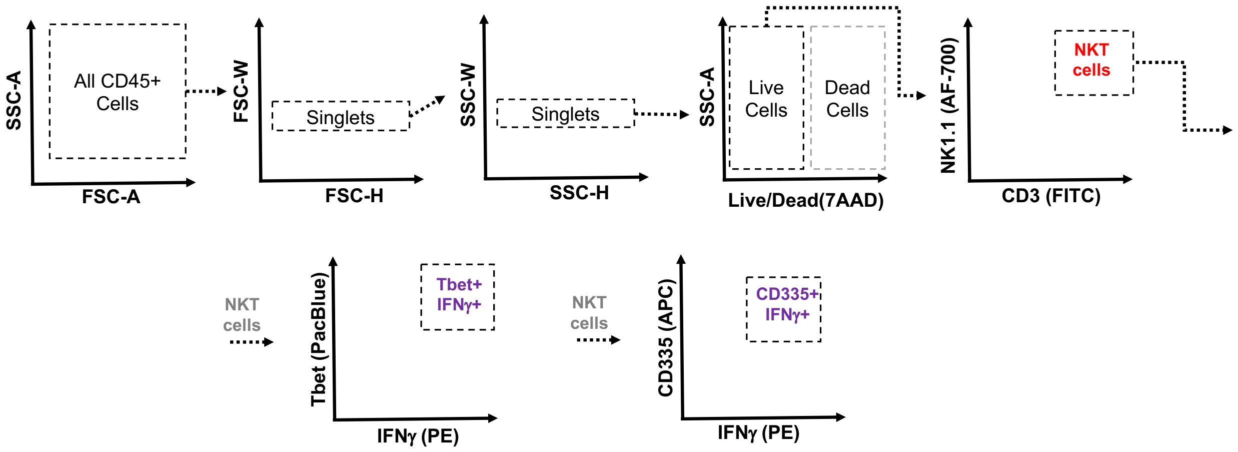


d) CD1d+ MECs analysis strategy:

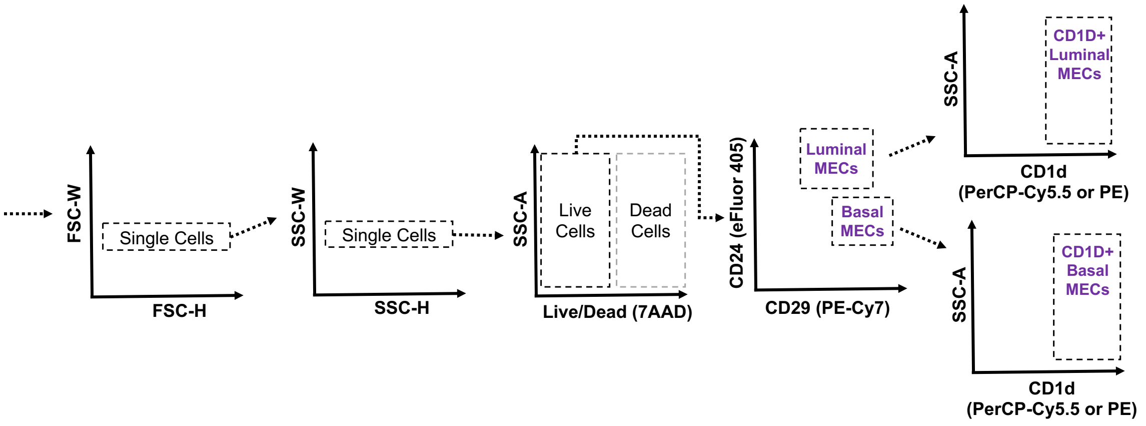

e) MHC-I and MHC-II MECs analysis strategy:

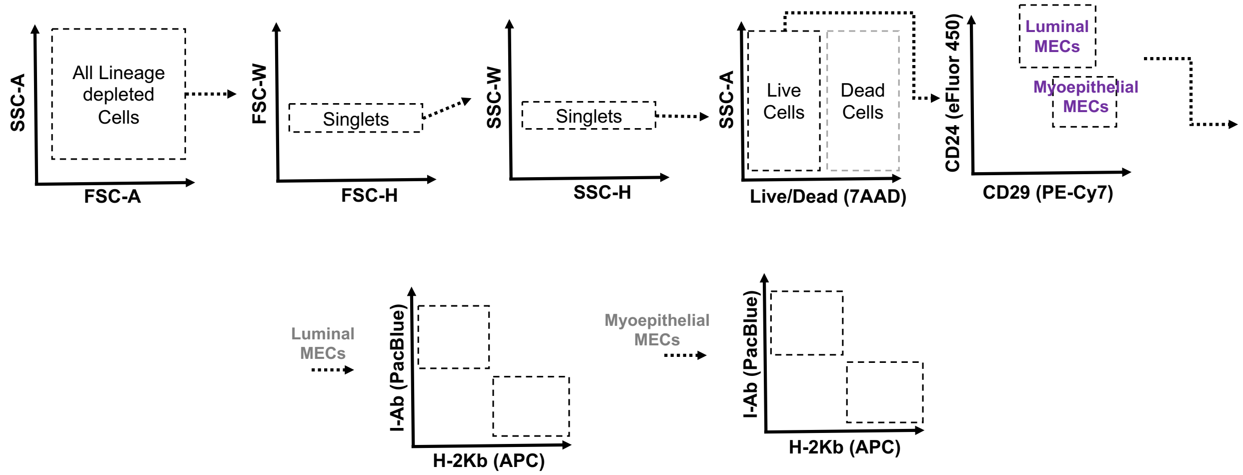


**Mammary Organoid Culture.** Mammary tissue dissected was minced and digested for ~40 minutes in Collagenase A, type IV solution (Sigma, #C5138-1G), following a series of centrifugations to enrich for mammary organoids. Freshly isolated mammary organoids were cultured with Essential medium (Advanced DMEM/F12, supplemented with ITS (Insulin/Transferrin/Sodium selenite, Gibco, #41400-045, and FGF-2 (PeproTech, #450-33)) prior to analysis. For experiments shown in Supplementary Fig. S7, organoid cultures were derived from normal mammary tissue from pre- or post-pregnancy Balb/C female mice (RRID:IMSR_CRL:028), cultured in the presence of FGF-2 for 6 days, following FGF-2 withdrawal for 24 hrs and then incubated with Complete medium (AdDf+++, supplemented with ITS (Final Concentration:1x, Insulin/Transferrin/Sodium Selenite, Gibco, #41400-045), 17-β-Estradiol (Final concentration: 40ng/mL, Sigma #E2758), Progesterone (Final concentration: 120ng/mL, Sigma #P8783), Prolactin (Final concentration: 120ng/mL ,Sigma #L4021), as previously described (Ciccone et al., 2020). For experiments shown in Supplementary Fig. S8, organoids cultures were derived from pre- or post-pregnancy CAGMYC MECs, following treatment with doxycycline (DOX, 0.1mg/mL) for 2 days (DD2). For experiments shown in Supplementary Fig. S12, organoid cultures were derived from NOD/SCID female mice, transplanted with either pre- or post-pregnancy CAGMYC MECs, following treatment with doxycycline (DOX, 0.1mg/mL) for 2 days (DD2).

**RT-qPCR.** Lineage depleted MECs or organoid cultures were washed with 0.5mL 1x PBS, following RNA extraction with Trizol (0.5mL, Thermo Fisher Scientific, #15596018). Reverse transcription was carried out using SuperScript III ™ kit (Thermo Fisher Scientific, #18080-051). RTqPCR was performed using a Quantstudio 6 with SYBR Green Master mix (Applied Biosystems, #4368577). Relative mRNA expression of target gene was calculated via the ΔΔCt method and normalized to *β-actin* mRNA levels.

Cd1d qPCR primers: FWD: 5’ TCC GGT GAC TCT TCC TTA CA 3’ and REV: 5’ CTG GCT GCT CTT CAC TTC TT 3’.

β-actin qPCR primers: FWD: 5’ TGT TAC CAA CTG GGA CGA CA 3’ and, REV: 5’ GGG GTG TTG AAG GTC TCA AA 3’.

**Mammary fat pad transplantation.** MaSCs-enrichment was performed as previously described (dos Santos et al., 2013). In short, lineage depleted MECs were incubated with biotinylated anti-CD1d antibody, to allow for MaSC enrichment. CD1d-enriched MEC fractions were resuspended with 50% growth factor reduced matrigel solution (Corning, #356230) and injected into the cleared fat-pad of the inguinal mammary gland (anterior part of the gland). For experiments presented on Supplementary Fig. S8 CD1d-enriched MECs fractions (~100K) were injected into the mammary fatpad of 12 weeks old CAG-only female mice, followed by DOX-treatment and histology analysis. For experiments presented on Supplementary Fig. S12 CD1d-enriched MECs fractions (~100K) were injected into the mammary fatpad of 12 weeks old NOD/SCID (RRID:IMSR_JAX:001303) female mice, followed by DOX-treatment and histology analysis. For experiments presented on Figure 5 and Supplementary Fig. S13, pre- or post-pregnancy CD1d WT CAGMYC MECs (~10K) or CD1d KO CAGMYC MECs (~10k) were injected into the mammary fatpad of 8-10 weeks old CD1d WT female mice, and allowed 3-days of tissue engraftment prior to DOX-treatment for 5 days.

**Histological analysis:** For histological analysis, the left inguinal mammary gland was harvested and fixed in 4% Paraformaldehyde overnight prior to paraffin embedding. For conventional histological analysis, mammary gland tissue slides were stained with Hematoxylin and Eosin (H&E). For ductal quantification, mammary gland H&E histological images were uploaded into Fiji (Fiji, RRID:SCR_002285), and ducts present in the posterior part of the gland were manually counted. Immunohistochemistry staining (IHC) was performed on a Roche Discovery Ultra Automated IHC/ISH stainer. For Masson's trichrome staining, Leica Multistainer Stainer/Coverslipper Combo (ST5020-CV5030) was used to stain slides according to standard reagents and protocols. Images were acquired using Aperio ePathology (Leica Biosystems) slide scanner in 40X lenses.

**Immunofluorescence (IF) analysis.** Paraffin-embedded mammary gland sections were deparaffinized in Xylene (Sigma Cat#534056) and rehydrated, followed by antigen retrieval in Trilogy (Cell Marque Cat# 920P-10). Tissue was washed in 1x PBS (phosphate-buffered saline) for 1 min then blocked with blocking solution (10mM Tris-HCl pH 7.4, 100mM $MgCl_2$, 0.5% Tween 20, 10% FBS, 5% goat serum) for 4 hours in a humidified chamber. Sections were stained with the appropriate conjugated primary antibodies in blocking solution for 16 hours at 4°C. After subsequent washings with 1x PBS and blocking solution, tissues were incubated with DAPI (Sigma Cat# 10236276001) for 10 minutes to stain nuclei, and slides were mounted in ProLong Glass Antifade Mountant (Invitrogen Cat# P36980). Cell visualization and image collection was performed on a Zeiss LSM780 confocal laser-scanning microscope utilizing Zen lite software, Blue edition (ZEN Digital Imaging for Light Microscopy, RRID:SCR_013672) version 2.0.0.0. Non-specific staining was defined as follows:

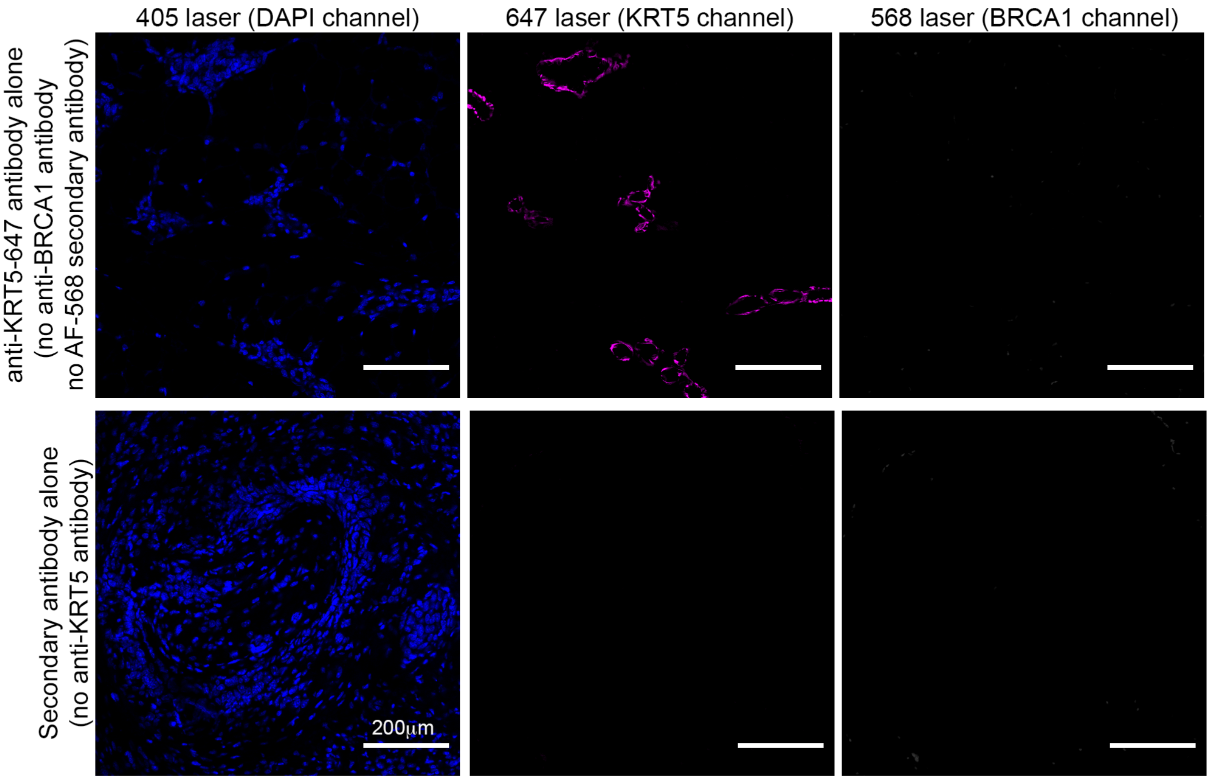


**Doxycycline (DOX) treatment.** Doxycycline was purchased from Takara Bio USA, Inc. (#631311) and sucrose was purchased from Sigma (S7903). DOX drinking solution (1 mg/mL) was prepared using sterile 1% sucrose water.

**Tamoxifen (TAM) treatment.** Tamoxifen USP grade was purchased from Sigma-Aldrich (Cat# 1643306) and sunflower seed oil (European Pharmacopoeia grade) was purchased from Sigma-Aldrich (Cat# 88921). To prepare the working solution, the Tamoxifen powder was weighed and dissolved in ethanol by vortexing. Heat sterilized sunflower oil was added at a ratio of 19:1 oil:ethanol mixture to a final concentration of 5mg/100ul (one dose), heated to 55°C and shaken vigorously to homogenize the mixture. $Krt5^{CRE-ERT2}Brca1^{fl/fl}p53^{het}$ transgenic female mice received a total of three intraperitoneal doses of Tamoxifen warmed to 37°C on alternate days.

**Monitoring tumor growth.** 3 week old $Krt5^{CRE-ERT2}Brca1^{fl/fl}p53^{-/+}$ female mice were treated with TAM. Half of TAM-treated female mice were housed together (pre-pregnancy/nulliparous group), and the other half were paired with a male (1 female and 1 male per breeding cage). Breeding TAM-treated females were allowed to give birth, nurse the offspring (21 days), and were considered post-pregnant (parous) after 40 days from offspring weaning. Both pre- and post-pregnancy mice were monitored for signs of tumor growth, and added to the Kaplan-Meier curve as soon as there was a palpable tumor. Mice with a tumor burden exceeding the limit of the animal's well-being (>2 cm), or mice showing signs of distress independently of tumor development were euthanized. At experimental end point, mammary tissue or mammary tumors were harvested for histological and flow cytometry analysis. Statistical analysis was performed with Logrank (Mantel-Cox) test.

**Western blot.** DOX-treated and control organoid cultures were homogenized in 1x Laemmli sample buffer (Bio-Rad, #1610747). Samples were loaded into home made 10% SDS-PAGE gel and transferred overnight to PVDF membrane (Bio-Rad, #162-0177) using wet-transfer apparatus. Membranes were blocked with 1% BSA solution and incubated overnight with a diluted solution of primary antibody, followed by incubation with HRP-conjugated antibody for 40 minutes. HRP signal was developed with Luminata Crescendo Western HRP substrate (Millipore, WBLUR0100) in autoradiography film (Lab Scientific, #XARALF2025). Developed films were scanned on Epson Perfection 2450 photo scanner.

**scRNA-seq data analysis.** Single cell data (pre-pregnancy mammary glands= 3,439 cells; post-pregnancy mammary glands= 4,412 cells) were aligned to mm10 using CellRanger v.3.1.0 (10x Genomics) (Cell Ranger , RRID:SCR_017344) (Zheng et al., 2017), and downstream processing was performed using Seurat v3.1.1 (SEURAT, RRID:SCR_007322) (Stuart et al., 2019). Cells with fewer than 250 features or higher than 10% mitochondrial gene content were removed prior to further analysis. Genes with fewer than 3 cells expressing them were removed, and the data were then log-normalized. Post-filtering analysis was performed on 3,075 cells (pre-pregnancy) and 4,029 cells (post-pregnancy). Principal component analysis was performed using the top 2,000 variable genes. This analysis was used to identify the number of significant components before clustering. Clustering was performed by calculating a shared nearest neighbor graph, using a resolution of 0.6. Subsetting into different cell types was performed using known markers for MECs, T-cells, Myeloid cells, B cells and NK cells. Epithelial cells for both datasets were defined by the expression of Epcam, Krt8, Krt18, Krt5 and Krt14(cluster average expression > 2). Non-epithelial were cells considered having low expression of Epcam, Krt8, Krt18, Krt5 and Krt14. Epithelial lineage identification and T-cell lineage identification was performed utilizing a previously validated gene signature (Henry et al., 2021). Genes used to define each immune cluster (differentially expressed genes, DEGs) were determined using known cell type markers and using the FindAllMarkers function, which uses a Wilcoxon Rank Sum test to identify differentially expressed genes between all clusters in the dataset. Cell cycle scoring was performed with the CellCycleScoring function, using the default gene lists provided by Seurat. Cell dendrograms were generated using the BuildClusterTree function in Seurat, using default arguments. Diffusion mapping was performed using the DiffusionMap function from the "destiny" R package (Angerer et al., 2016). Gene Set Enrichment Analysis (GSEA, RRID:SCR_003199) (Subramanian et al., 2005) was used for global analyses of differentially expressed genes.

**RNA-seq library preparation and analysis.** FACS-isolated pre- and post-pregnancy NKT cells were collected and homogenized in TRIzol LS (Thermo Fisher Scientific, #10296010) for RNA extraction. Double stranded cDNA synthesis and Illumina libraries were prepared utilizing the Ovation RNA-seq system (V2) (Nugen Technologies, #7102-32). RNA-seq libraries were prepared utilizing the Ovation ultralow DR multiplex system (Nugen Technologies, #0331-32). Each library (n=2 per experimental condition) was barcoded with Illumina TruSeq adaptors to allow sample multiplexing, followed by sequencing on an Illumina NextSeq500, 76bp single-end run. Analyses were performed with command-line interfaced tools such as FastQC (FastQC, RRID:SCR_014583) (Andrews, 2015) for quality control and Trimmomatic (Trimmomatic, RRID:SCR_011848) (Bolger et al., 2014) for sequence trimming. We used STAR (STAR, RRID:SCR_004463) for mapping reads (Dobin et al., 2013), FeatureCounts (featureCounts, RRID:SCR_012919) for assigning reads to genomic features (Liao et al., 2014) and DESeq (DESeq, RRID:SCR_000154) to assess changes in expression levels simultaneously across multiple conditions and in multi-factor experimental designs, incorporating information from multiple replicates (Anders and Huber, 2010). Genes with a statistically significant pvalue of $p < 0.05$ were considered differentially expressed. Gene Set Enrichment Analysis (GSEA) (Gene Set Enrichment Analysis, RRID:SCR_003199) was used for global analyses of differentially expressed genes (Subramanian et al., 2005). GSEA terms with statistically significant pvalue of $p<0.05$ were selected for data plotting and data interpretation. For experiments presented on Figure 2D, FACS-isolated, pre- and post-pregnancy CD45+NK1.1+CD3+ NKT cells (n=2 females per experimental group, n=4 pairs of mammary glands per female, n=2 biological replicates per experimental group) were utilized. For experiments presented on Supplementary Figure S12, total mammary tissue isolated from DOX-treated, NOD/SCID female mice transplanted with either pre- or post-pregnancy CAGMYC MECs (n=2 biological replicates per group) were utilized.

**ChIP-seq library analysis.** Previously published H3K27ac ChiP-seq datasets (Feigman et al., 2020) were mapped to the indexed mm9 genome using bowtie2 short-read aligner tool (Langmead et al., 2009), using default settings. MACS2 peak-calling program (MACS, RRID:SCR_013291) (Zhang et al., 2008) was used to identify enriched genomic regions in this data by comparing the pulldown ChIP data to the control (Input) data using a q-value cutoff of $1.00^{-3}$ . Identification of genes closest to these differentially called peaks was performed using Genomic Regions Enrichment of Annotations Tool (UCSC Genome Browser, RRID:SCR_005780) (McLean et al., 2010). Peak visualizations were generated using the UCSC Genome Browser (UCSC Genome Browser, RRID:SCR_005780) (Dreszer et al., 2012).

**Cut&Run library analysis.** Previously published H3K27ac Cut&Run datasets (Ciccone et al., 2020), were mapped to the indexed mm9 genome using bowtie2 short-read aligner tool (Langmead et al., 2009) using default settings. Sparse Enrichment Analysis for Cut&Run (SEACR) peak-calling program (Meers et al., 2019) was used to identify enriched genomic regions with an empirical threshold of n=0.01, returning the top n fraction of peaks based on total signal within peaks. The stringent argument was implemented, which used the summit of each curve. Identification of genes closest to these differentially called peaks was performed using Genomic Regions Enrichment of Annotations Tool (UCSC Genome Browser, RRID:SCR_005780) (McLean et al., 2010). Peak visualizations were generated using the UCSC Genome Browser (UCSC Genome Browser, RRID:SCR_005780) (Dreszer et al., 2012).

**ATAC-seq library preparation and analysis.** Nuclei of FACS-isolated, pre- and post-pregnancy NKT cells were isolated utilizing hypotonic lysis buffer and incubated with Tn5 enzyme from Nextera DNA sample Preparation kit (Illumina, #FC-121-1031) for the preparation of ATAC libraries. Each library (n=2 per experimental condition) was amplified and barcoded as previously described (Buenrostro et al., 2013), then pooled for sequencing on an Illumina Nextseq500, 76bp single-end run. ATACseq library reads (n=2 per cell condition) were mapped to the indexed mm9 genome using Bowtie2 short read-aligner (Bowtie 2, RRID:SCR_016368) (Langmead et al., 2009) and replicate alignment files were merged. MACS2 (MACS, RRID:SCR_013291) (Zhang et al., 2008) was used to identify enriched genomic regions in both conditions using a tag size of 25bp and a q-value cutoff of $1.00^{-2}$. Peaks were annotated using Homer (HOMER, RRID:SCR_010881) (Benner et al., 2017) with standard mm9 genome reference. Location of peaks was then grouped into intergenic, promoter and genic (containing 5'UTR, Exons, Introns, Transcription Termination Sites, 3'UTR, ncRNA, miRNA, snoRNA, and rRNA) regions. The UCSC genome browser (UCSC Genome Browser, RRID:SCR_005780) (Dreszer et al., 2012) was used to analyze genomic regions for overlap, using the Bedtools intersect function (BEDTools, RRID:SCR_006646) (Quinlan and Hall, 2010) . Any base pair overlap was enough to consider two regions "shared" and regions where no overlap existed defined the regions as exclusively being in one condition. The comparison was made into a venn-diagram using tool available at https://www.meta-chart.com/venn.

**DNA motif analysis.** Peaks from pre- and post-pregnancy NKT cells ATAC-seq libraries were utilized as input for for an unbiased transcription factor analyses using Analysis of Motif Enrichment (AME) (McLeay and Bailey, 2010) and Find Individual Motif Occurrences (FIMO) (MEME Suite - Motif-based sequence analysis tools, RRID:SCR_001783) (Grant et al., 2011) was used to computationally define DNA binding motif regions to identify sequences of known motifs, with a statistical threshold of 0.0001.

**Genomic library preparation and Copy number variation (CNV) analysis.** Mammary normal tissue and tumor from nulliparous BRCA1 KO p53het female mice were dissociated as above described. Lineage depleted tumor cells were utilized for DNA extraction using DNeasy Blood & Tissue Kit (Qiagen 69504). Genomic DNA was sonicated to an average of 300 bp using Covaris E220 Focused-ultrasonicator. For library preparation, fragmented DNA went through standard end-repair (NEB E6050), dA-tailing (NEB E6053), and sequencing adapter ligation (NEB M2200) steps. Following universal adapter ligation, eight cycles of PCR was performed for each sample. During the PCR step, a unique pair of Illumina TruSeq i7 index and i5 index was added to each sample. The PCR library was purified with AMPure XP beads (Beckman Coulter A63881), and quantified using NanoDrop spectrophotometer and Agilent Technologies 2100 Bioanalyzer. Whole-genome-sequencing libraries with different combination of Illumina indexes were pooled together for one lane of Illumina MiSeq. 150 base pairs from both ends were sequenced along with two 8-bp indexes. For CNV analysis, Read 1 of the sequence data was mapped to the mm9 reference genome using Hisat2 version 2.1.0 in single read alignment mode (Kim et al., 2015). The reference genome was divided into 5,000 variable-length bins with equal mappability as previously described (Baslan et al., 2012). The ratio of mapped reads in the tumor sample to mapped reads in the diploid sample (normal tissue) was used to compute a fitted piecewise constant function (segmentation). This segmentation used DNAcopy version 1.50.1 implementation of the circular binary segmentation algorithm (Seshan and Olshen, 2014) and the copy number profiles were plotted using R version 3.4.4 (R Core Team, 2019).

#### Supplementary Figures

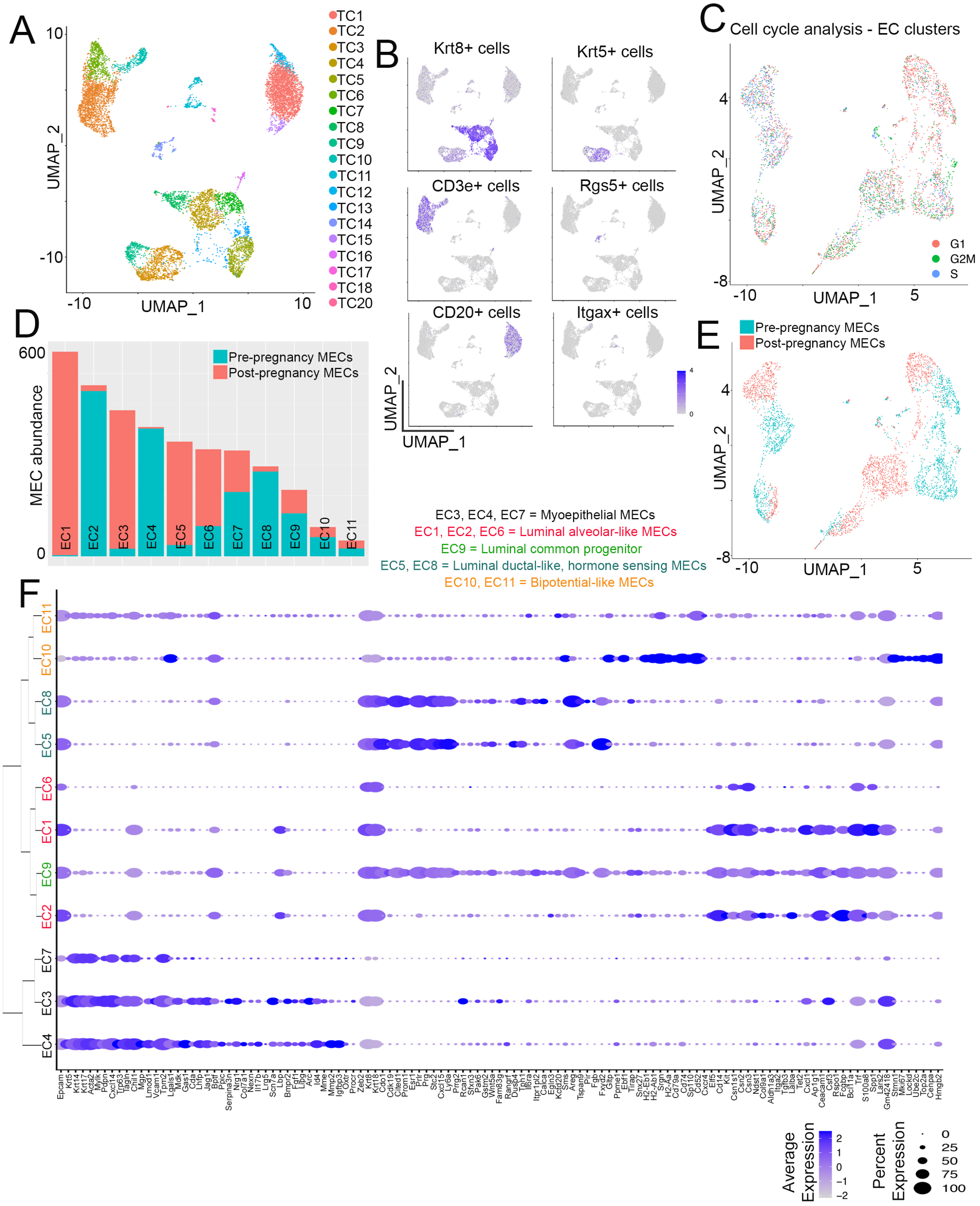


### Supplementary Figure S1

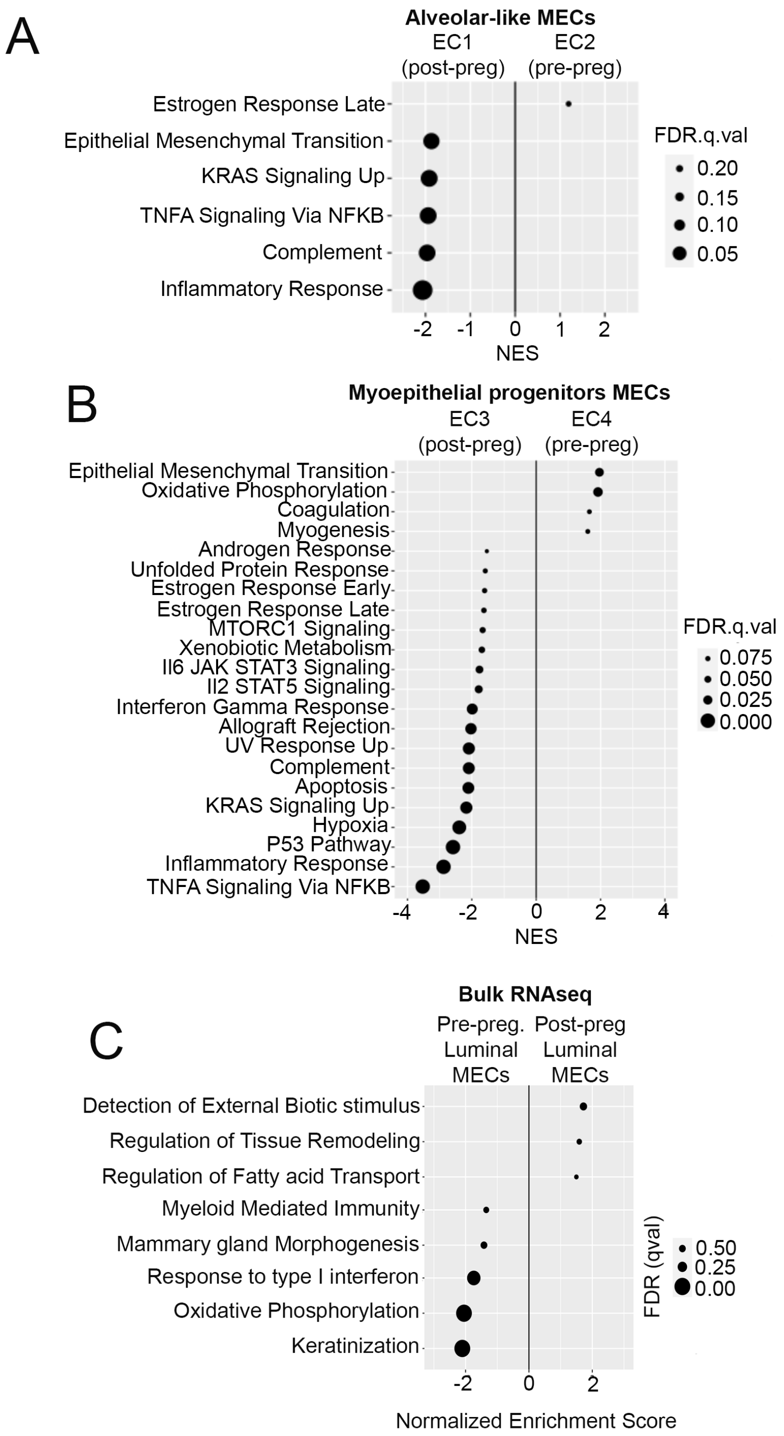


324 Supplementary Figure S2

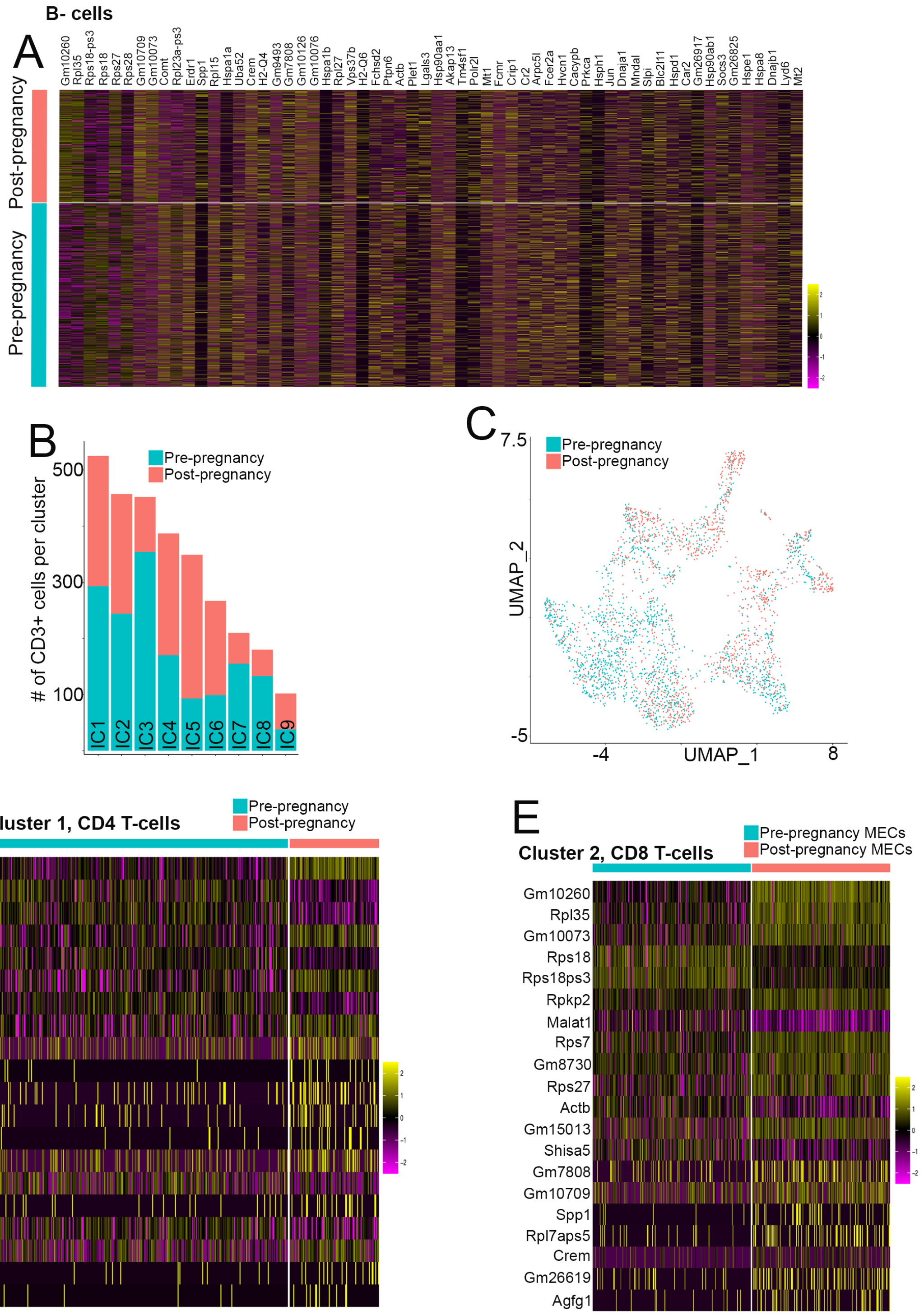


Supplementary Figure S3

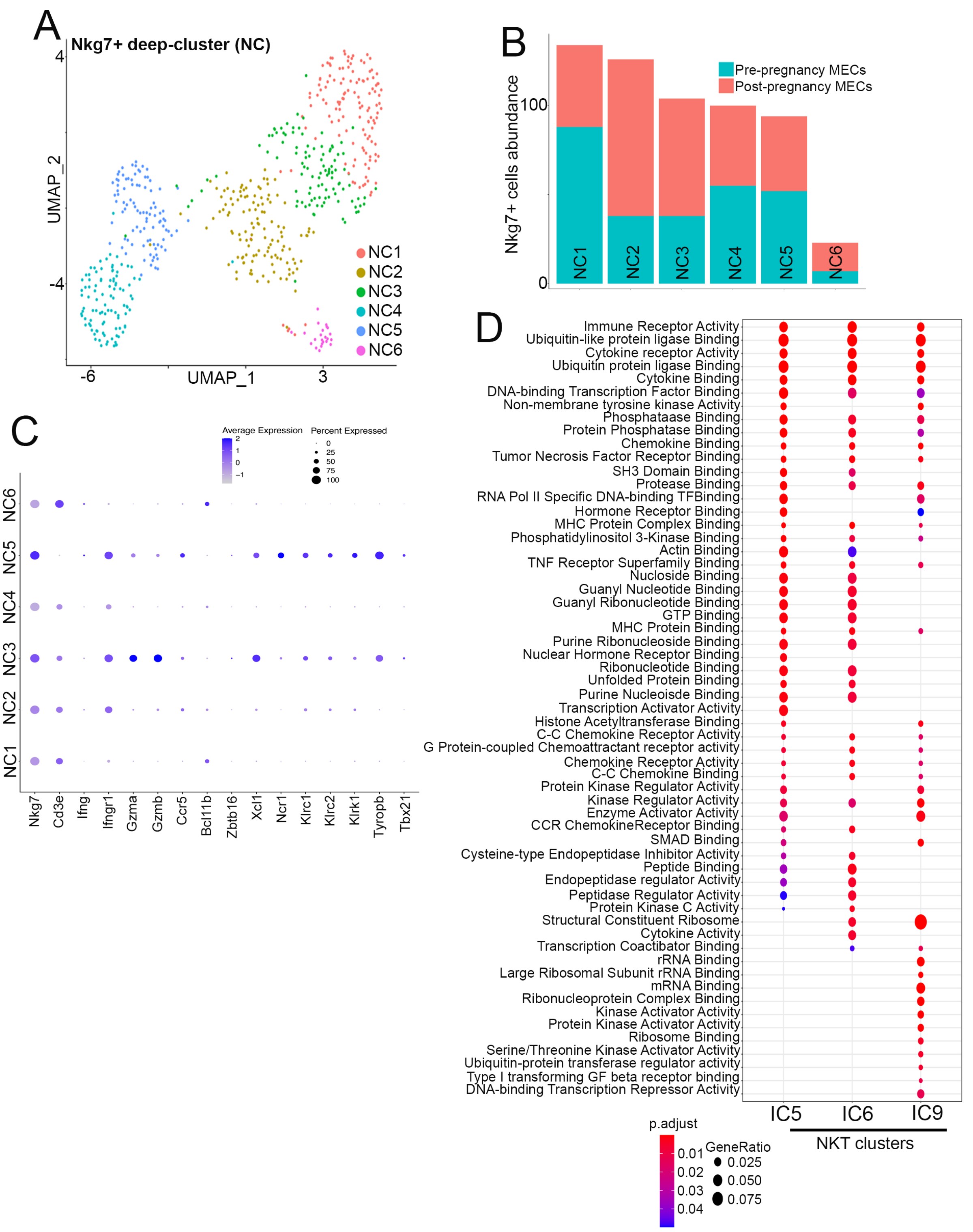


326 Supplementary Figure S4

327

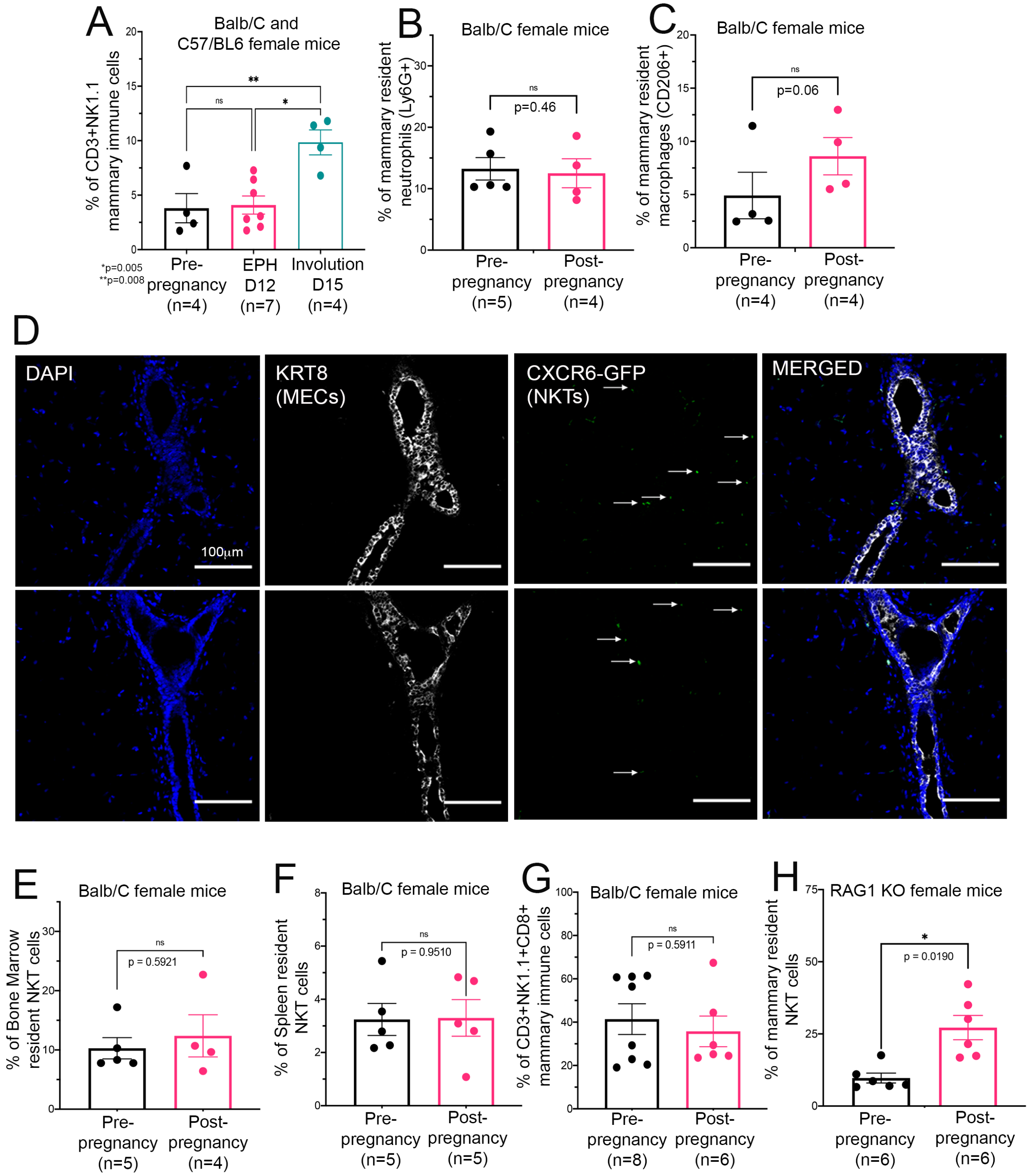


328 Supplementary Figure S5

A

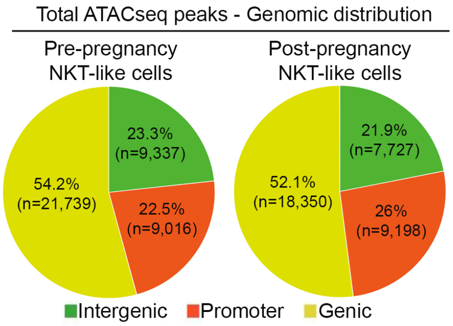


B

| Transcription factor | DNA Motif | Pre-pregnancy NKT cells | Post-pregnancy NKT cells |
|---|---|---|---|
| T-bet (Tbx21) | 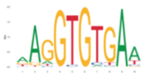 | 11.2% of total ATAC-seq peaks | 10.7% of total ATAC-seq peaks |
| Plzf (Zbtb16) | 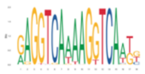 | 11.2% of total ATAC-seq peaks | 10.7% of total ATAC-seq peaks |
| Egr2 | 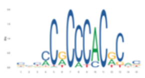 | 47.3% of total ATAC-seq peaks | 40.5% of total ATAC-seq peaks |

C

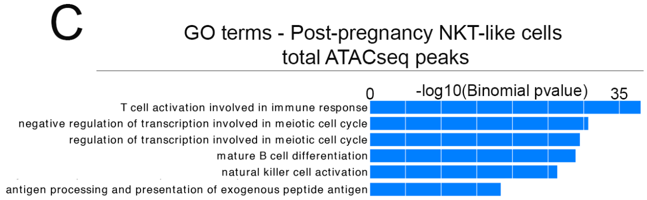


D

| Transcription factor | DNA Motif | Pre-pregnancy NKT cells DNA motif enrichment (p-value) | Post-pregnancy NKT cells DNA motif enrichment (p-value) |
|---|---|---|---|
| NFATC2 | 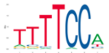 | 9.52 E-25 | 7.22 E-32 |
| BACH2 | 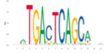 | 3.84 E-15 | 2.99 E-26 |
| FOSL1 | 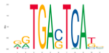 | 2.24 E-02 | 1.36 E-11 |
| JUNB | 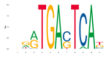 | 2.14 E-02 | 2.87 E-11 |
| MAF::NFE2 | 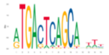 | 1.47 E-02 | 7.67 E-11 |

Supplementary Figure S6

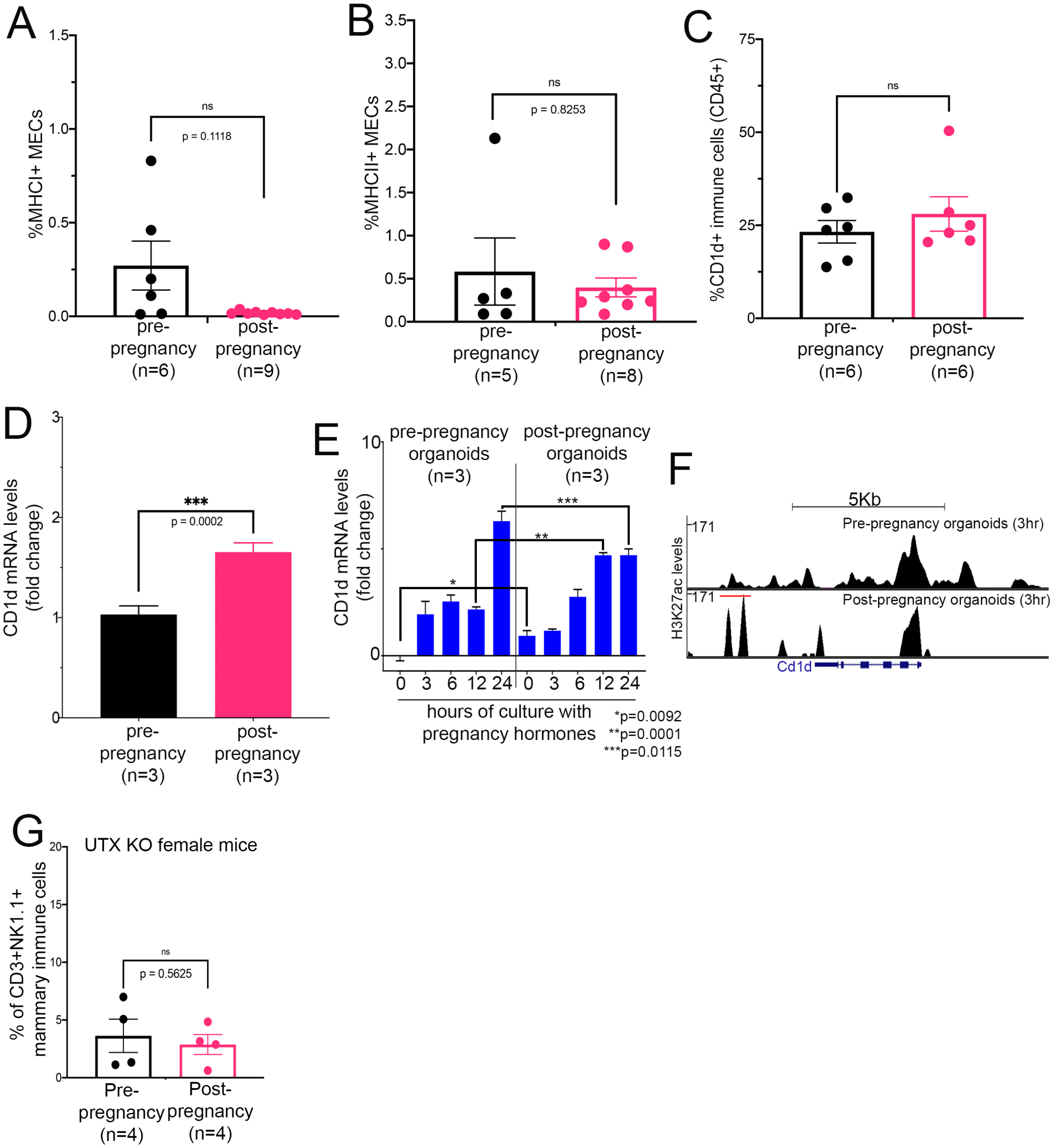


### Supplementary Figure S7

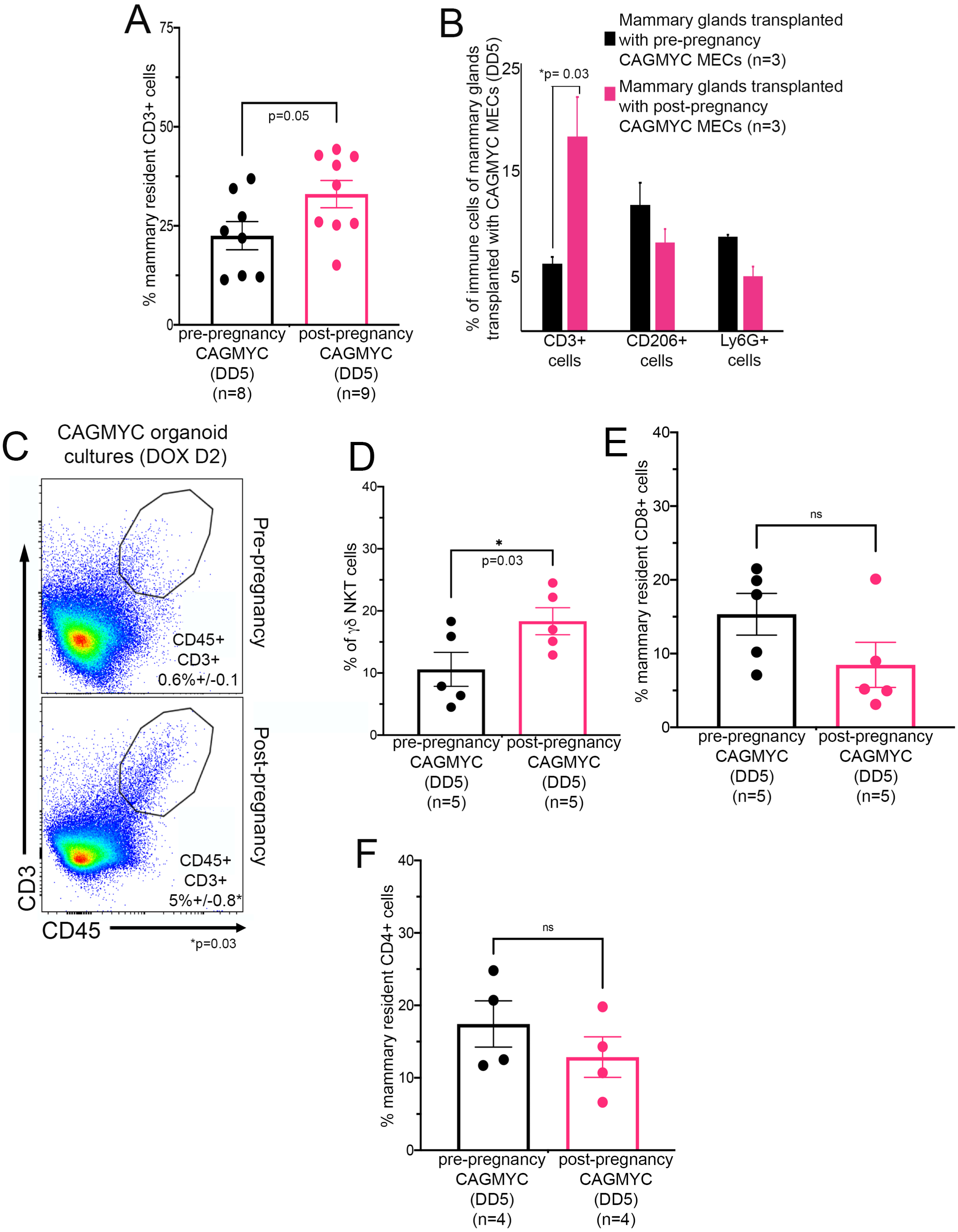


Supplementary Figure S8

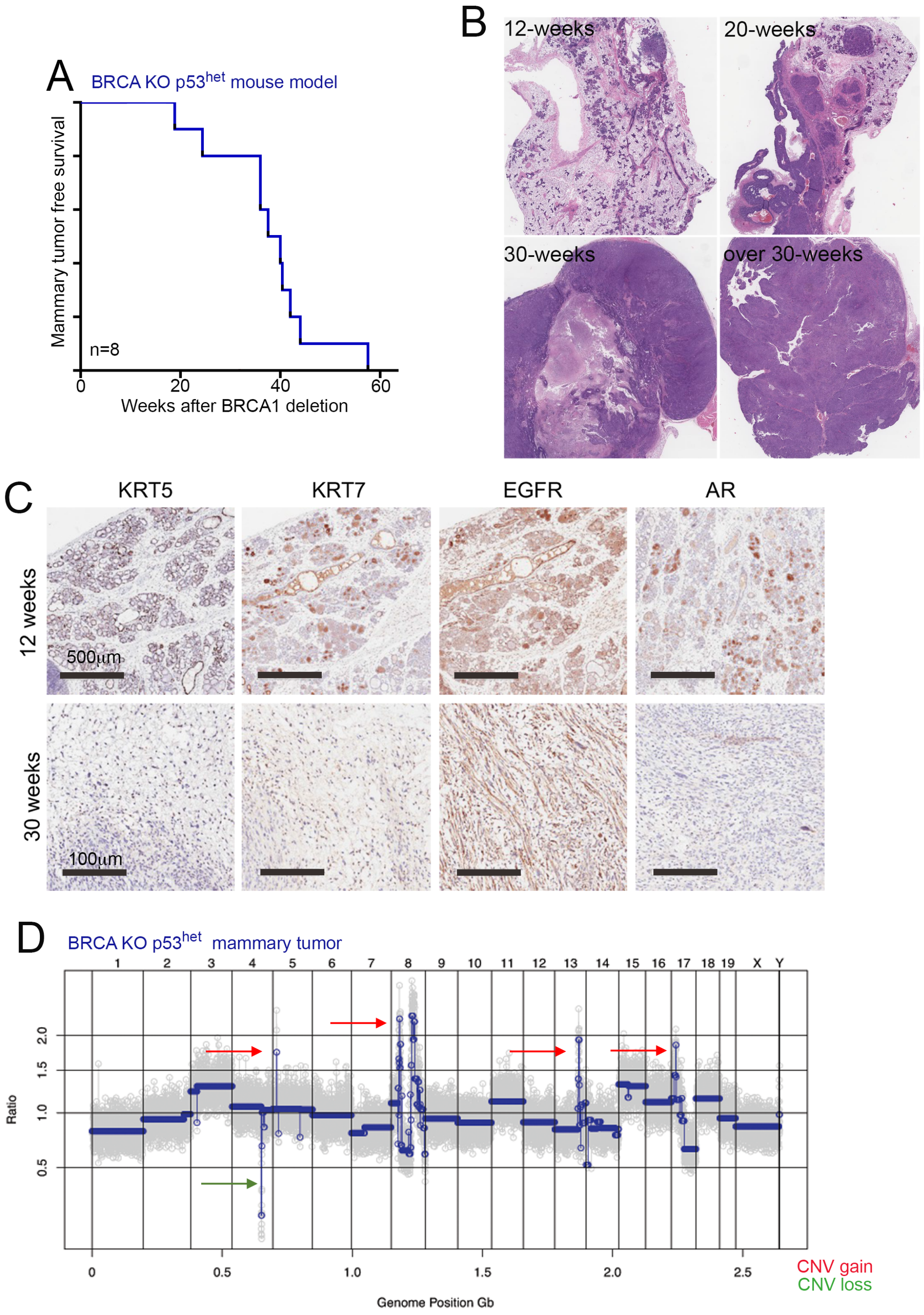


### Supplementary Figure S9

334

335

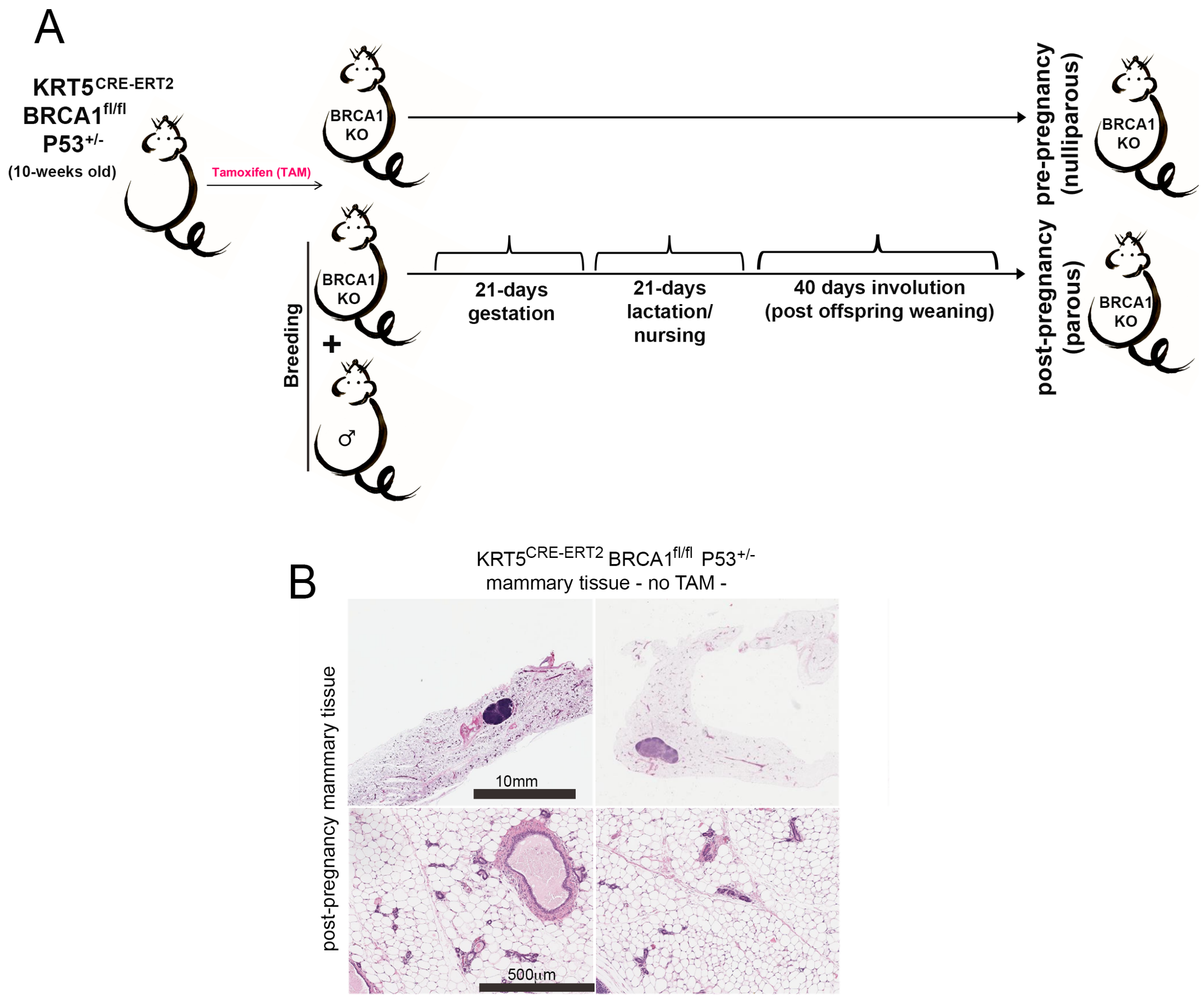


Supplementary Figure S10

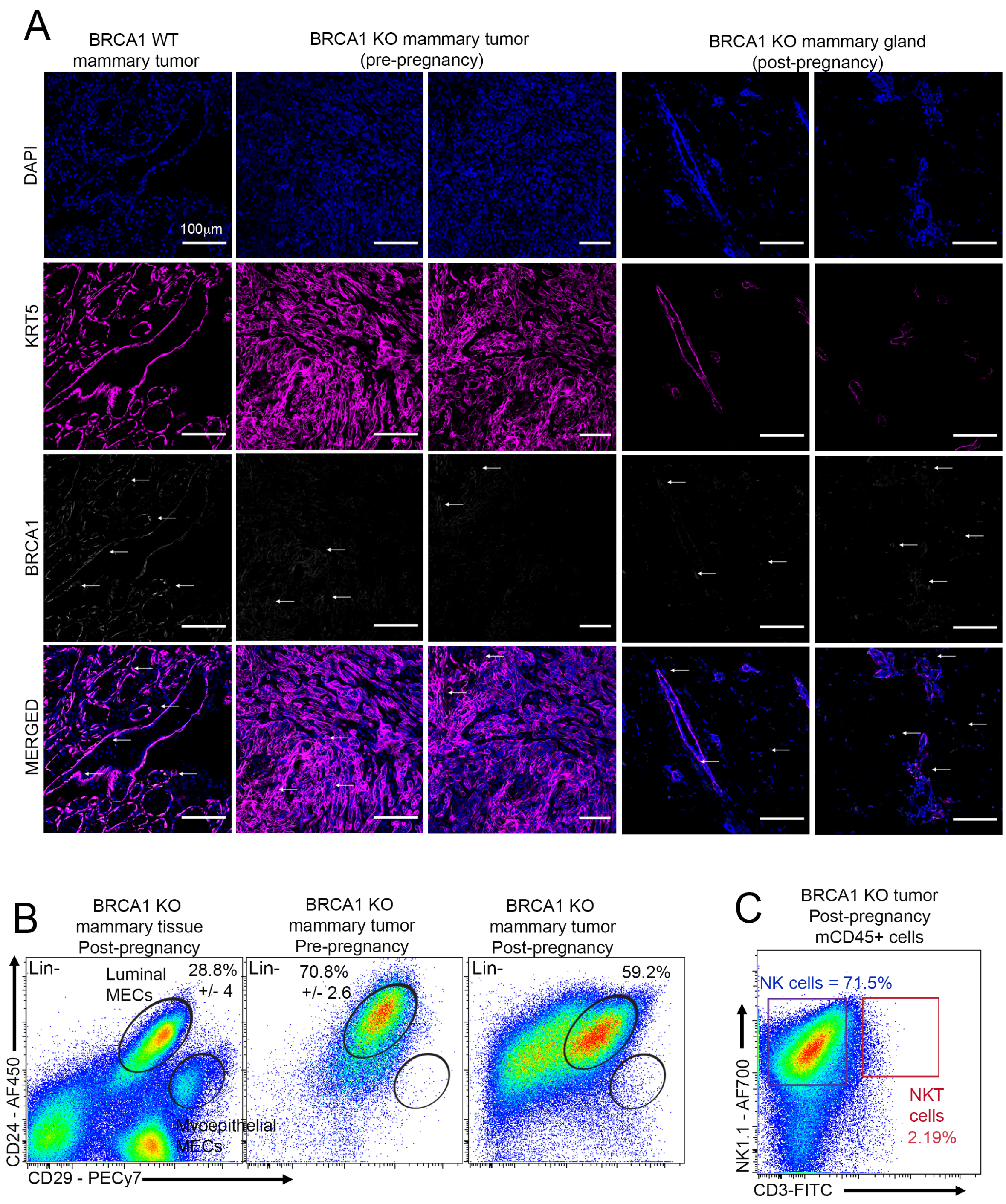


Supplementary Figure S11

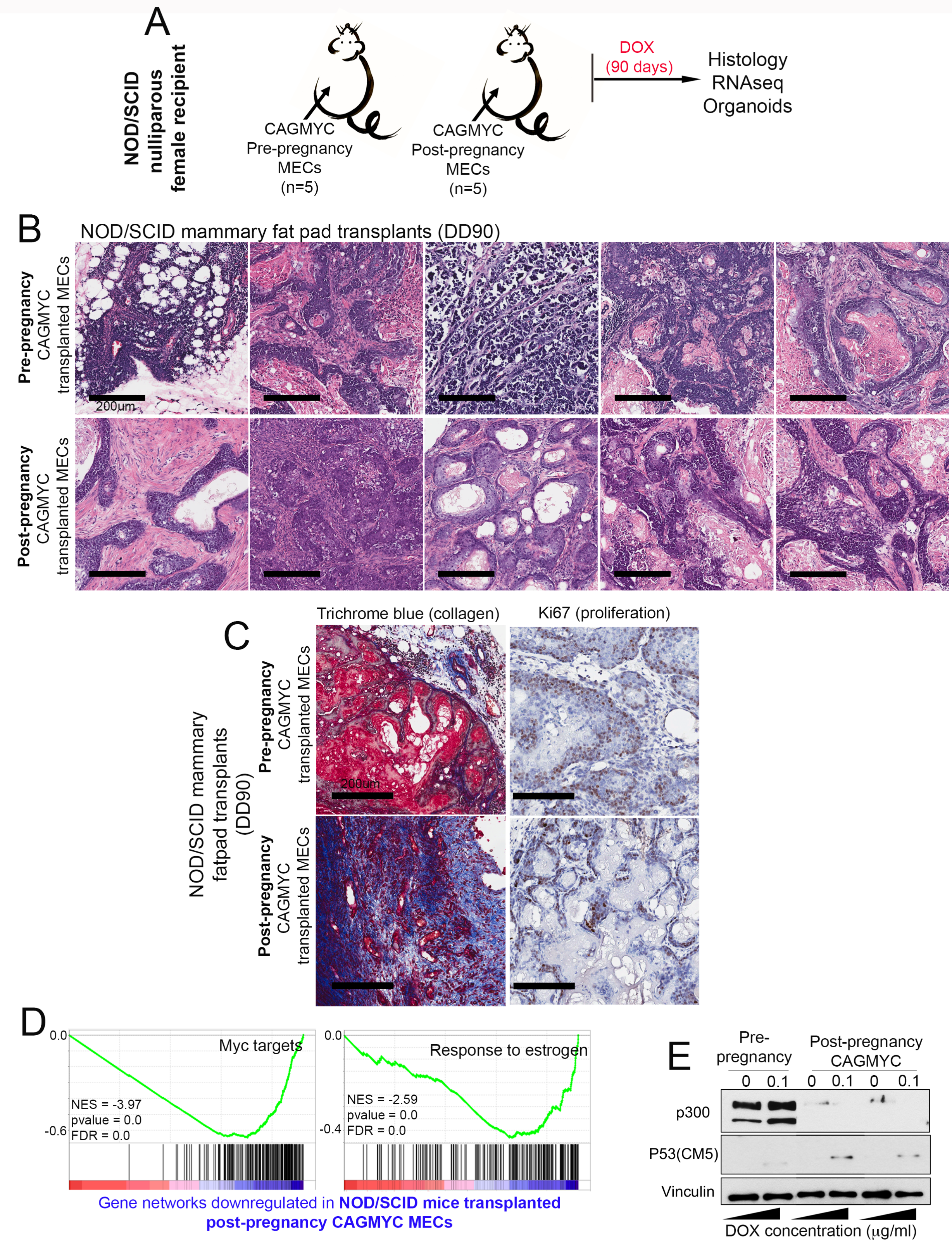


### Supplementary Figure S12

340

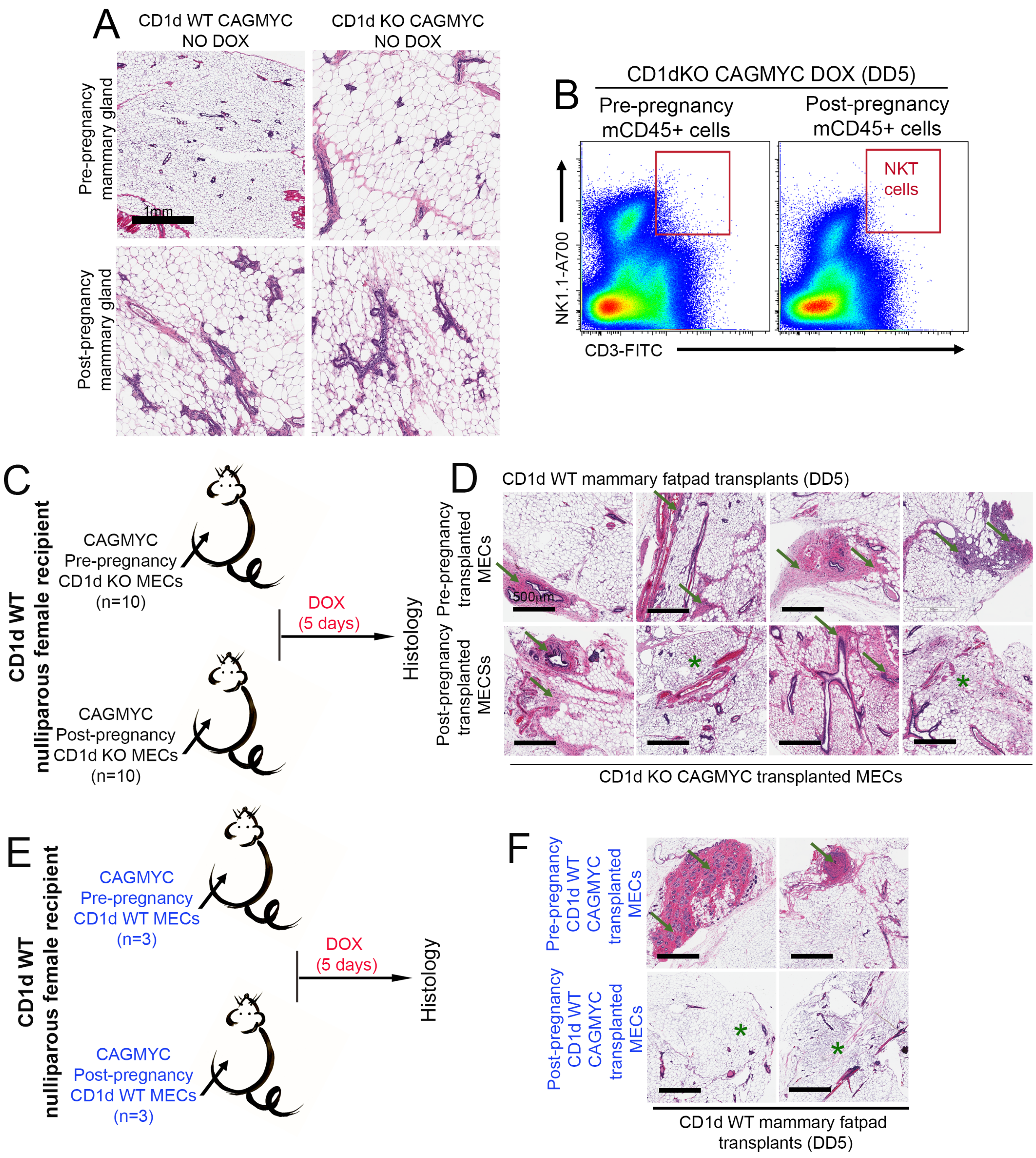


Supplementary Figure S13

341

342

**Supplementary Figures Legend**

**Figure S1. Single cell levels classification of pre- and post-pregnancy MECs.** (**A**) UMAP showing the distribution of total, pre- and post-pregnancy mammary resident cells.(**B**) Classification of mammary resident lineages based on the expression of *Krt8* and *Krt5* (epithelial cells), *Cd20* (B-cells), *Cd3e* (T-cell), *Rags5* (Fibroblasts), and *Itgax* (myeloid cells). (**C**) UMAP showing the distribution of mammary epithelial clusters according different stages of cell cycle. **D**) Cell abundancy of pre- and post-pregnancy epithelial cells across all 11 epithelial clusters. **(E**) UMAP showing epithelial-focused re-clustering (Epcam+, Krt8+,Krt18+, and Krt5+ cells) of pre-pregnancy (blue) and post-pregnancy (pink) MECs. (**E**) dot plot analysis of molecular signatures and lineage identity of pre- and post-pregnancy MECs.

**Figure S2. Pathway analysis post-pregnancy biased epithelial cells.** (**A, B**) Gene set enrichment analysis (GSEA) of pathways differentially enriched in (A) alveolar-like MECs, and (B) myoepithelial progenitor-like MECs. (**C**) Gene set enrichment analysis (GSEA) of pathways differentially enriched in FACS-isolated, pre- and post-pregnancy luminal MECs.

**Figure S3. Characterization of pre- and post-pregnancy mammary resident immune cells.** (**A**) Heatmap showing top DEGs across pre- and post-pregnancy B-cell clusters. (**B**) Cell abundancy of pre- and post-pregnancy CD3+ immune cells. (**C**) UMAP showing CD3+ focused re-clustering of pre-pregnancy (blue) and post-pregnancy (pink) immune resident cells. (**D-E**) Heatmap showing top 20 DEGs across CD4+ T-cells (**D**, cluster 1) and CD8+ T-cells (**E**, cluster 2) harvested from pre and post-pregnancy mammary tissue.

**Figure S4. scRNA-seq identification of post-pregnancy NKT cells.** (**A**) UMAP showing Nkg7+ expressing cells focused re-clustering (NKT/NK cells) of pre- and post-pregnancy mammary tissue. (**B**) Cell abundancy of pre- and post-pregnancy Nkg7+ immune cells. (**C**) Dot plot showing the expression of NKT/NK associated genes across Nkg7+ cell clusters. (**D**) Dotplot of pathway analysis across populations of CD3+ NKT cells (IC5, IC6 and IC9).

**Figure S5. Cellular characterization of post-pregnancy mammary immune microenviroment.** (**A**) Quantification of NKT cells abundance in mammary tissue from nulliparous female mice (black bar, n=4), from female mice during Exposure to Pregnancy Hormone day 12 (pink bar, EPH D12, n=7), and from female mice at post-pregnancy involution D15 (blue bar, n=4). EPH D12 x Involution D15 *p=0.005; Involution D15 x Pre-pregnancy **p=0.008. (**B**) Quantification of Ly6G+ mammary resident neutrophils abundance in tissue from pre- and post-pregnancy female mice. n=5 nulliparous and n=4 parous female mice. *p=0.46. (**C**) Quantification of CD206+ mammary resident macrophages abundance in tissue from pre- and post-pregnancy female mice. n=4 nulliparous and n=4 parous female mice. *p=0.06. (**D**) Immunofluorescence images (IF) of mammary tissue from Cxcr6-GFP-KI nulliparous mouse model showing the presence of GFP+ cells (NKT cells, green, white arrows) surrounding duct structures (Krt8+, white). (**E**) Quantification of NKT cells abundance in bone marrow from pre- and post-pregnancy female mice. n=5 nulliparous and n=4 parous female mice. *p=0.5921. (**F**) Quantification of NKT cells abundance in spleen from pre- and post-pregnancy female mice. n=5 nulliparous and n=5 parous female mice. *p=0.95. (**G**) Quantification of CD8+ NKT cells abundance in mammary tissue from pre- and post-pregnancy female mice. n=8 nulliparous and n=6 parous female mice. *p=0.6 (**H**) Quantification of NKT cells abundance in mammary tissue from pre- and post-pregnancy RAG1 KO female mice. n=6 nulliparous and n=6 parous female mice. *p=0.019. For all analyses, error bars indicate standard error of mean across samples of same experimental group. Statistically significant differences were considered with Student's t-test *p*-value lower than 0.05 (p<0.05).

**Figure S6. The molecular signature of post-pregnancy NKT cells.** (**A**) Genomic distribution of total ATAC-seq peaks from FACS-isolated pre- and post-pregnancy NKT cells. (**B**) TF motif analysis across total ATAC-seq peaks from FACS-isolated pre- and post-pregnancy NKT cells. (**C**) GO term analysis of genes associated with total ATAC-seq peaks of FACS-isolated post-pregnancy NKT cells. (**D**) TF motif analysis across exclusive ATAC-seq peaks from FACS-isolated pre- or post-pregnancy NKT cells.

**Figure S7. The effects of pregnancy in controlling CD1d expression.** (**A**) Quantification of MHC-I+ MECs in pre- and post-pregnancy mammary tissue. n=6 nulliparous and n=9 parous female mice. *p=0.1. (**B**) Quantification of MHC-II+MECs in pre- and post-pregnancy mammary tissue. n=5 nulliparous and n=8 parous female mice. *p=0.8. (**C**) Quantification of CD1d+ immune cells in pre- and post-pregnancy mammary tissue. n=6 nulliparous and n=6 parous female mice. *p=0.28. (**D**) qPCR analysis of *Cd1d* mRNA levels in lineage depleted pre- and post-pregnancy MECs. n=3 biological replicates. p=0.0002. (**E**) qPCR analysis of *Cd1d* mRNA levels in organoid cultures derived from pre- and post-pregnancy MECs treated with pregnancy hormones. n=3 biological replicates. 0h, *p=0.0092; 12h, **p=0.0001; 24h, ***p=0.01 (**F**) Genome browser tracks showing distribution of SEACR-called, H3K27ac Cut&Run, peaks at the Cd1d genomic locus in organoid cultures derived from pre- and post-pregnancy MECs treated for 3 hours with pregnancy hormones (**G**) Quantification of NKT cells abundance in mammary tissue from nulliparous and parous UTX KO female mice, which are deficient for activated NKt CELLS. n=4 nulliparous and n=4 parous female mice. *p=0.5. For all analyses, error bars indicate standard error of mean across samples of same experimental group. Statistically significant differences were considered with Student's t-test *p*-value lower than 0.05 ($p<0.05$).

**Figure S8. The effects of cMYC-overexpression on pregnancy-induced immune changes and CD1d+ post-pregnancy MECs.** (**A**) Quantification of CD3+ immune cell abundance in mammary tissue from DOX-treated (DD5), pre- and post-pregnancy CAGMYC female mice. n=8 nulliparous and n=9 parous female mice. p=0.05. (**B**) Quantification of CD3+ (T/NKT cells), CD206+ (macrophages) and Ly6G+ (neutrophils) in mammary tissue of DOX-treated nulliparous female mice transplanted with pre-pregnancy CAGMYC MECS (black bars, n=3) or post-pregnancy CAGMYC MECs (pink bar, n=3) *p=0.03. (**C**) Quantification of CD45+CD3+ immune cell abundance in DOX-treated (2-days), organoid cultures derived from pre-pregnancy CAGMYC mammary tissue (top panel, n=3) or from post-pregnancy CAGMYC mammary tissue (bottom panel, n=3) *p=0.03. (**D**) Quantification of $\gamma\delta$TCR expression at the surface of NKT cells in mammary tissue from DOX-treated (DD5), pre- and post-pregnancy CAGMYC female mice. n=5 nulliparous and n=5 parous female mice. *p=0.03. (**E**) Quantification of total CD8+ T-cells in mammary tissue from DOX-treated (DD5), pre- and post-pregnancy CAGMYC female mice. n=5 nulliparous and n=5 parous female mice. *p=0.24. (**F**) Quantification of total CD4+ T-cells in mammary tissue from DOX-treated (DD5), pre- and post-pregnancy CAGMYC female mice. n=4 nulliparous and n=4 parous female mice. *p=0.41. Error bars indicate standard error of mean across samples. Statistically significant differences were considered with Student's t-test *p*-value lower than 0.05 ($p<0.05$).

**Figure S9. Characterization of Krt5$^{CRE-ERT2}$Brca1$^{ko}$p53$^{het}$ (Brca1 KO) mouse model.** (**A**) Tumor-free survival plot of nulliparous Brca1 KO female mice in weeks after TAM-treatment (to induce Brca1 deletion) (n=8). (**B**) Mammary tissue and tumors from Brca1 KO nulliparous female mice at specific time points after TAM-treatment. (**C**) Immunohistochemistry (IHC) analysis of mammary tissue and tumors from Brca1 KO nulliparous female mice for marker of basal-like mammary tumors KRT5, KRT7, EGFR and AR. (**D**) Genomic segment plot showing Copy Number Variation (CNV) in mammary tumor from nulliparous Brca1 KO female mice.

**Figure S10. Probing the role of pregnancy in Brca1 KO mammary tumor development.** (**A**) Experimental approach showing the strategy for Brca1 deletion, and analysis of tumor development in pre- and post-pregnancy Brca1 KO female mice. (**B**) H&E histological images of normal mammary tissue harvested from parous, no TAM treated, $Krt5^{CRE-ERT2}Brca1^{fl/fl}p53^{het}$ (therefore Brca1 WT) female mice.

**Figure S11. Tissue and cellular analysis of post-pregnancy Brca1 KO mammary tissue.** (**A**) Immunofluorescence analysis (IF) of BRCA1 protein levels (white signal) in mammary epithelial cells (KRT5+, pink signal) from Brca1 WT mammary tumor (far left panels), pre-pregnancy Brca1 KO mammary tumors (left and middle panels), and from normal mammary tissue from parous Brca1 KO female mice (right and far right panels). Arrows indicate cells positive for BRCA1 and KRT5. (**B**) FACS plots showing the abundance of luminal mammary epithelial MECs ($CD24^{high}CD29^{low}$) and myoepithelial mammary epithelial MECs ($CD24^{low}CD29^{high}$) in mammary tissue from parous Brca1 KO female mice (left panel), mammary tumor from nulliparous Brca1 female mice (middle panel), and mammary tumor from parous Brca1 female mice (right panel). (**C**) FACS plots showing the abundance of NKT cells in mammary tumors from parous Brca1 KO female mice.

**Figure S12. cMYC-overexpression induces oncogenesis of post-pregnancy MECs transplanted into NOD/SCID mammary fatpads.** (**A**) Experimental approach showing strategy for the transplantation of pre- and post-pregnancy CAGMYC MECs into the fatpad of nulliparous NOD/SCID female mice, and tissue analysis. (**B**) H&E stained histology images from DOX-treated (DD90) NOD/SCID mammary tissue transplanted with pre- and post-pregnancy CAGMYC MECs. n=5 nulliparous and 5 parous female mice. Scale: 200μm. (**C**) Immunohistochemistry (IHC) analysis of Masson's Trichrome levels (collagen deposition) and Ki67 levels (proliferation) in DOX-treated (DD90) NOD/SCID mammary tissue transplanted with pre- and post-pregnancy CAGMYC MECs. n=5 nulliparous and 5 parous female mice. Scale: 200μm. (**D**) Gene set enrichment analysis of gene networks down-regulated in NOD/SCID mice transplanted with post-pregnancy CAGMYC MECs. (**E**) Western blot of p300, and p53 proteins in organoid cultures derived from NOD/SCID transplanted pre- and post-pregnancy CAGMYC tumor cells, with and without DOX treatment (2 days). Vinculin protein levels were used as endogenous control.

**Figure S13. Loss of CD1d expression supports the malignant transformation and oncogenesis of post-pregnancy CAGMYC MECs.** (**A**) H&E stained histology images from mammary tissue from pre- and post-pregnancy CD1d WT CAGMYC and CD1d KO CAGMYC female mice without DOX treatment. Scale: 1mm. (**B**) Flow cytometry analysis of mammary resident NKT cells harvested from pre- and post-pregnancy CD1d KO CAGMYC female mice (DOX D5). (**C**) Experimental approach showing the strategy for the transplantation of pre- and post-pregnancy CD1d KO CAGMYC MECs into the fatpad of nulliparous CD1d WT female mice, and tissue analysis. (**D**) H&E stained histology images from mammary tissue from DOX-treated (DD5) CD1d WT nulliparous female mice transplanted with pre- and post-pregnancy CD1d KO CAGMYC MECs Scale: 500μm. (**E**) Experimental approach showing the strategy for the transplantation of pre- and post-pregnancy CD1d WT CAGMYC MECs into the fatpad of nulliparous CD1d WT female mice, and tissue analysis. (**F**) H&E stained histology images from mammary tissue from DOX-treated (DD5) CD1d WT nulliparous female mice transplanted with pre- and post-pregnancy CD1d WT CAGMYC MECs Scale: 500μm. Green arrows indicate signs of malignant lesions/mammary hyperplasia. Green asterisks indicate normal-like ductal structures.
